# Full-length isoform transcriptome of developing human brain provides new insights into autism

**DOI:** 10.1101/2020.06.27.175489

**Authors:** Kevin Chau, Pan Zhang, Jorge Urresti, Megha Amar, Akula Bala Pramod, Jiaye Chen, Amy Thomas, Roser Corominas, Guan Ning Lin, Lilia M. Iakoucheva

## Abstract

Alternative splicing plays important role in brain development, however its global contribution to human neurodevelopmental diseases (NDD) has not been fully investigated. Here, we examined the relationships between full-length splicing isoforms expression in the brain and *de novo* loss-of-function mutations identified in the patients with NDDs. We analyzed the full-length isoform transcriptome of the developing human brain and observed differentially expressed isoforms and isoform co-expression modules undetectable by gene-level analyses. These isoforms were enriched in loss-of-function mutations and microexons, co-expressed with a unique set of partners, and had higher prenatal expression. We experimentally tested the impact of splice site mutations in five NDD risk genes, including *SCN2A*, *DYRK1A* and *BTRC,* and demonstrated exon skipping. Furthermore, our results suggest that the splice site mutation in *BTRC* reduces translational efficiency, likely impacting Wnt signaling through impaired degradation of β-catenin. We propose that functional effect of mutations associated with human diseases should be investigated at the isoform-rather than the gene-level resolution.

**Highlights:**

- Differential isoform expression analysis of the human brain transcriptome reveals neurodevelopmental processes and pathways undetectable by differential gene expression analyses.
- Splicing isoforms impacted by neurodevelopmental disease (NDD) risk mutations exhibit higher prenatal expression, are enriched in microexons and are involved in neuronal-related functions.
- Isoform co-expression network analysis identifies modules with splicing and synaptic functions that are enriched in NDD mutations.
- Splice site mutations impacting NDD risk genes cause exon skipping and produce novel isoforms with altered biological properties.
- Functional impact of mutations should be investigated at the full-length isoform-level rather than the gene-level resolution

## Introduction

More than 95% of multi-exon human genes undergo alternative splicing (AS) and/or use alternative promoters to increase transcriptomic and proteomic diversity, with an estimated average of five to seven isoforms transcribed per gene (Pan et al., 2008; Steijger et al., 2013; Wang et al., 2008). Alternative splicing is highly specific, and expression of isoforms is often restricted to certain organs, tissues or cell types (Barbosa-Morais et al., 2012; Sapkota et al., 2019; Shalek et al., 2013; Trapnell et al., 2010). In addition, many isoforms are expressed only during specific developmental periods (Kalsotra and Cooper, 2011). The alternatively spliced isoforms encoded by the same gene can also be expressed at different levels in the same tissue or during the same developmental period (Wang et al., 2008).

The developing human brain exhibits one of the highest frequencies of alternative splicing events (Calarco et al., 2011; Mele et al., 2015; Raj and Blencowe, 2015; Yeo et al., 2004). Many of the processes occurring during neural development including cell-fate determination, neuronal migration, axon guidance and synaptogenesis, are controlled by differentially expressed alternatively spliced isoforms (Grabowski, 2011; Kim et al., 2013; Li et al., 2007). Several recent studies, including one by us, began to investigate isoform-level transcriptome dysregulation in psychiatric diseases (Gandal et al., 2018; Li et al., 2018; Parikshak et al., 2016). However, spatio-temporal analyses of the full-length isoform transcriptome of the developing human brain remains relatively unexplored.

Integration of the brain spatiotemporal transcriptome with the genetic data from exome and whole genome sequencing studies have provided important insights into neurodevelopmental diseases (NDDs) (Li et al., 2018; Lin et al., 2015; Parikshak et al., 2013; Satterstrom et al., 2020; Willsey et al., 2013). Most of the recent work in this area has been focused on understanding the effect of mutations at the gene-level resolution, whereas the isoform-specific impact of loss-of-function (LoF) mutations in the context of brain development has not yet been fully investigated.

It is important to map LoF mutations to transcripts, because protein isoforms encoded by different transcripts have drastically different protein interaction capabilities. As we have previously demonstrated, the majority of the isoforms encoded by the same gene share less than a half of their interacting partners in the human interactome network (Yang et al., 2016). This observation points to striking functional differences between splicing isoforms that are not accounted for by the majority of the existing gene-level studies. In addition, our recent work demonstrated that isoform-level networks of autism risk genes and copy number variants provide better resolution and depth around disease proteins (Corominas et al., 2014).

To better understand how NDD risk mutations dysregulate normal brain development, we analyzed the temporal isoform transcriptome of the developing human brain using the BrainSpan RNA-seq dataset from the PsychEncode Consortium (Li et al., 2018) summarized to full-length isoforms as previously described (Gandal et al., 2018). We identified hundreds of differentially expressed isoforms (DEI) and dozens of isoform co-expression modules at brain developmental periods starting from fetal to adult. When compared to the gene-level transcriptome, the full-length isoform transcriptome provides more meaningful insights and paints a more complete picture of neurodevelopmental processes. Importantly, many DEIs and isoform co-expression modules were undetectable by the gene-level analyses. Mapping autism spectrum disorder (ASD) risk mutations to DEI revealed that ASD LoF-impacted isoforms have higher prenatal expression, more frequently carry microexons, and are preferentially involved in key neuronal processes compared to non-impacted isoforms. Furthermore, isoform co-expression modules with splicing-related and synaptic functions were enriched in LoF-impacted isoforms implicating these functions in NDDs. Finally, we experimentally tested the impact of several splice site LoF mutations and demonstrated that they cause exon skipping to produce novel isoforms with altered biological properties. Our study makes a strong case for investigation of disease mutations at the full-length isoform-rather than the gene-level resolution.

## Results

### Construction, quality control, and validation of the full-length isoform transcriptome of the developing human brain

To investigate global patterns of isoform expression across brain development, we analyzed the temporal full-length isoform transcriptome of the developing human brain (**Supplementary Figure 1**). We used the BrainSpan RNA sequencing dataset from the PsychEncode Consortium (Li et al., 2018) summarized to full-length isoforms as previously described (Gandal et al., 2018). After rigorous quality control that included sample outlier detection through weighted gene co-expression analysis (WGCNA) (Oldham et al., 2012) and detection of confounding variables with surrogate variable analyses (Leek and Storey, 2007) (**Supplementary Figures 2-5, Materials and Methods**), we obtained expression profiles for 100,754 unique full-length isoforms corresponding to 26,307 brain-expressed human genes, resulting in ∼3.8 isoforms/gene.

To experimentally validate the quality of the BrainSpan isoform transcriptome, we used quantitative PCR (RT-qPCR) to estimate relative expression difference of 44 unique isoforms of 32 genes between two independent RNA samples that were age, sex and brain region-matched to the samples from the BrainSpan (**Materials and Methods**). The relative RT-qPCR isoform expression values in the independent samples of the frontal lobe of a 22-week-old fetus and the cerebral cortex of a 27-year-old adult were compared to the values obtained from BrainSpan. We observed positive correlation (R^2^=0.46) between experimental and BrainSpan-derived values for these isoforms, despite using independent samples for validation (**Supplementary Table 1, Supplementary Figure 6, Materials and Methods**). This correlation was comparable to our previous validation values from other PsychEncode datasets (R^2^=0.48 for ASD and R^2^=0.51 for SCZ) (Gandal et al., 2018).

### Differential isoform expression reveals distinct signals relative to differential gene expression

We recently demonstrated that isoform-level changes capture larger disease effects than gene-level changes in the context of three major psychiatric disorders (Gandal et al., 2018). Here, we investigated the role of full-length isoform expression in the context of the normal brain development. We performed differential expression analysis among all pairs of adjacent developmental periods as well as between pooled prenatal (P02-P07) and pooled postnatal (P08-P13) (PrePost) samples, yielding sets of differentially expressed genes (DEG) and DEI, respectively (**Materials and Methods**, **Supplementary Tables 2 and 3**). We observed the largest number of both DEGs and DEIs in the P06/P07 (late mid-fetal/late fetal) and P07/P08 (late fetal/neonatal) developmental periods, supporting critical brain remodeling right before and after the birth (**Fig. 1A**). In P06/P07, 8.3% of genes and 20.3% of isoforms (converted to gene identifiers) were differentially expressed, whereas in P07/P08 13.2% of genes and 20.4% of isoforms (converted to gene identifiers) were differentially expressed (**Supplementary Table 4**). Overall, 48.4% of genes and 64.9% of isoforms (converted to gene identifiers) were differentially expressed between prenatal and postnatal (PrePost) periods. These results indicate that expression levels of over half of all isoforms significantly change between prenatal and postnatal periods, suggesting profound transcriptomic remodeling during brain development.

**Figure 1.**
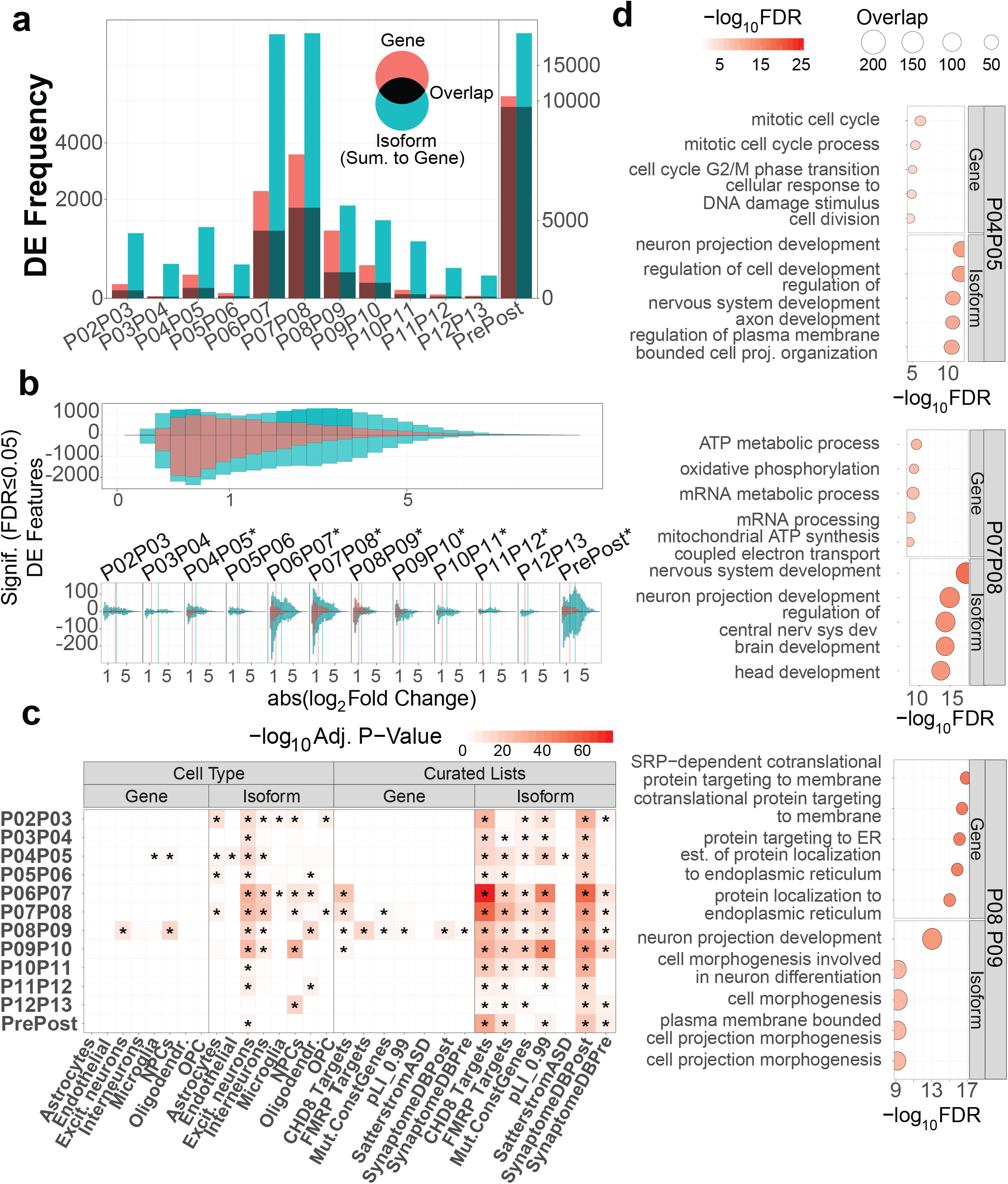
Differential gene and isoform expression analyses. **(A)** Number of significantly differentially expressed genes and isoforms in the adjacent brain developmental periods, and in prenatal vs postnatal periods. Isoform identifiers were summarized to gene identifiers for simplicity of comparison. Shaded areas represent identifiers shared between gene and isoform datasets, whereas unshaded bars represent genes (red) or isoforms (turquoise) unique to each dataset. (B) Effect size (absolute log2 fold change) distribution of differentially expressed genes (red) and isoforms (turquoise) of combined data (top) or per developmental period (bottom). Average absolute effect sizes for genes and isoforms are marked by corresponding colored vertical lines, and differences were tested using two-sample T-tests (*FDR < 0.05). **(C)** Enrichment of cell types and literature curated gene sets among genes and isoforms unique to each dataset (unshaded sets from a panel). Fisher’s exact test was used to calculate p-values. **(D)** Gene Ontology (GO) enrichment of differentially expressed genes and isoforms unique to each dataset (unshaded sets from a panel). Three adjacent periods are shown as examples (P04/05, P07/08 and P08/09). DEI are enriched in nervous system-related processes.

In addition to the greater fraction of DEI between adjacent and PrePost periods, we also observed significantly increased effect sizes (absolute log2 fold changes) among DEI as compared to DEG, both overall and in nearly every developmental period (**Fig. 1B**). This suggests that levels of differential expression were more pronounced at the full-length isoform level relative to the gene level, consistent with previous results obtained from NDD patient postmortem brains (Gandal et al., 2018). Thus, the full-length isoform transcriptome is likely to provide additional information about brain development that is missed by the gene transcriptome.

To better understand the biological basis of brain transcriptome differences at the gene and isoform levels, we performed enrichment analyses of unique non-overlapping DEGs and DEIs (lightly shaded subsets from **Fig. 1A**) using cell type and literature-curated gene lists (**Fig. 1C**). We used a published gene-level single cell RNA sequencing dataset (Zhong et al., 2018) for cell type enrichment, and NDD-related gene lists for gene enrichment analyses in each period as well as in the PrePost dataset (**Materials and Methods**). Overall, DEGs captured weaker enrichment signals than DEIs, potentially due to smaller DEG dataset sizes. Among cell types, DEIs were significantly enriched in excitatory neuron markers, especially in the prenatal to early childhood developmental periods (Fisher-exact test, max Bonferroni-adjusted P < 1E-09, OR = 2.39 – 3.29, min. 95% CI = 2.07, max 95% CI = 3.98 for P02/P03-P09/P10) (**Fig. 1C, left panel**). The DEIs from almost all periods were also enriched in post-synaptically expressed genes, as well as FMRP and CHD8 targets, with most significant enrichment during P06/P07 (late mid-fetal/late fetal). Interestingly, the DEIs from only P04/P05 (early mid-fetal) were enriched in autism risk genes (Satterstrom et al., 2020) (Fisher-exact test, Bonferroni-adjusted P = 0.005, OR = 3.88, 95% CI = 2.11 – 3.68) (**Fig. 1C, right panel**), and this signal was not observed at the gene level. The mid-to-late fetal developmental period was previously identified as critical to ASD pathogenesis (Parikshak et al., 2013; Willsey et al., 2013 Lin, 2015 #6310).

Functional Gene Ontology (GO) analyses in P04/P05, P07/P08 and P08/P09 demonstrated stronger enrichment of DEI in neurodevelopment-relevant processes compared to DEGs (**Fig. 1D, Supplementary Tables 5 and 6**). For example, “neuron projection development”, “brain development”, and “nervous system development” were enriched among DEIs, but not among DEGs. In contrast, DEGs were enriched in basic biological function-related processes, such as “mitotic cell cycle”, “metabolic processes”, “protein targeting” and “localization”. This suggests that the full-length isoform transcriptome provides better biological insights into brain development than the gene transcriptome.

### Differentially expressed isoforms impacted by autism loss-of-function mutations have higher prenatal expression

To improve understanding of the impact of NDD mutations on brain development, we mapped rare *de novo* loss-of-function (LoF) variants identified in the largest autism exome sequencing study (Satterstrom et al., 2020) to the full-length isoform transcriptome. A total of 12,111 ASD case variants and 3,588 control variants were processed through Ensembl’s Variant Effect Predictor (VEP) and filtered for consequences likely to result in the loss-of-function of the impacted gene or isoform (**Materials and Methods**, **Supplementary Table 7**). In total, 1,132 ASD case and 262 control variants fit this criterion, impacting 4,050 isoforms from 1,189 genes. At the isoform level, 3,128 isoforms were impacted by ASD case variants (ASD LoF), 848 isoforms by control variants (Control LoF), and 74 isoforms by both. We also defined a dataset of isoforms that were not impacted by ASD variants (Non-impacted by ASD LoF) as an internal control.

In every prenatal developmental period, as well as in the pooled prenatal sample, the expression of the ASD LoF-impacted isoforms was found to be significantly higher than Control LoF-impacted isoforms or Non-impacted by ASD LoF isoforms (Mann-Whitney test, BH-adjusted P-value ≤ 0.05) (**Fig. 2A**). This suggests that the potential decrease or loss of expression of these highly expressed isoforms in the normal prenatal human brain as a result of LoF mutation may contribute to ASD pathogenesis.

**Figure 2.**
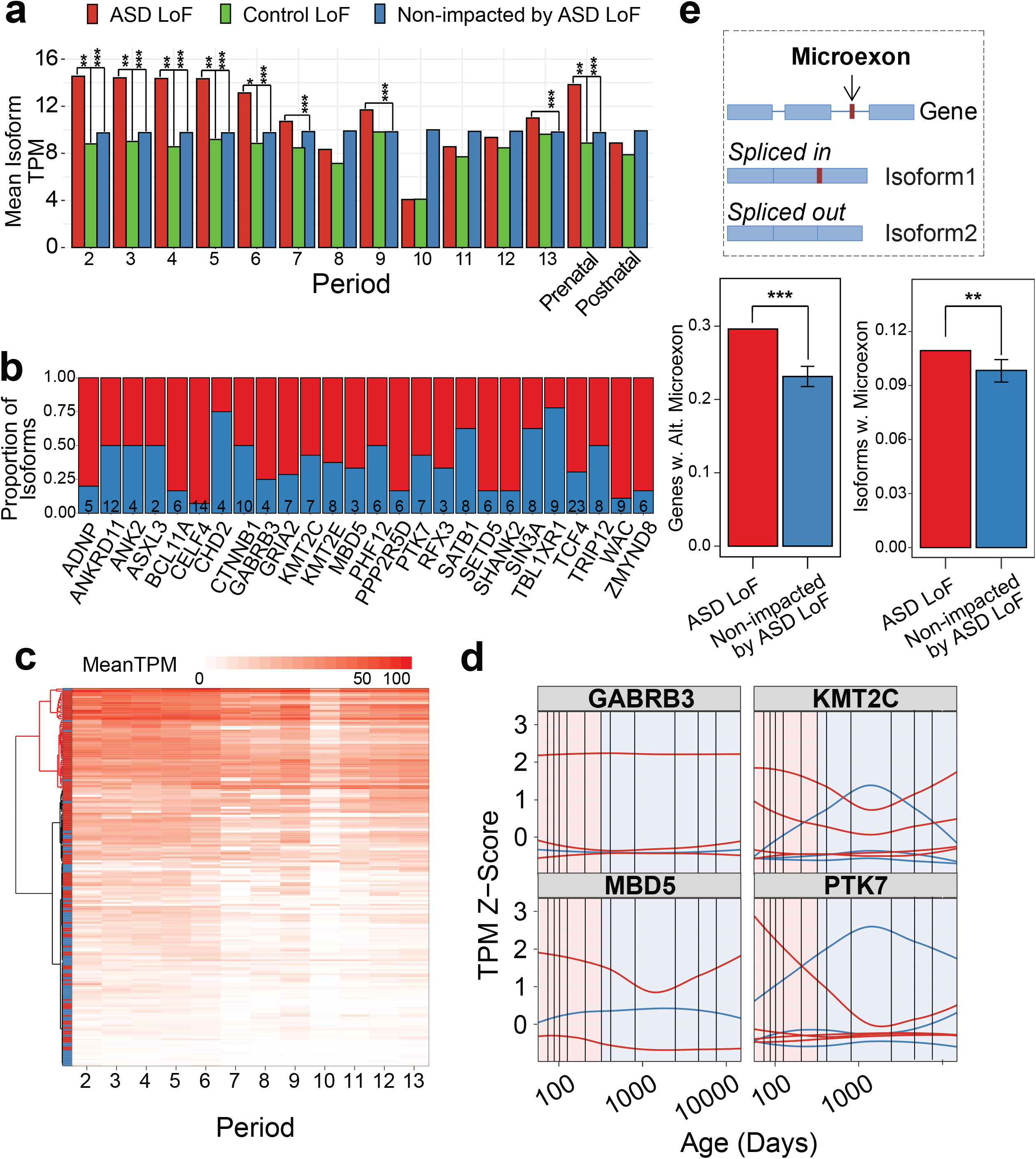
**Analyses of isoforms impacted by rare *de novo* ASD loss of function variants**. **(A)** Mean expression of isoforms impacted by case rare *de novo* ASD LoF variants (Impacted by ASD LoF) is significantly higher in prenatal periods compared to those impacted by control LoF (Impacted by Control LoF) mutations or to non-impacted isoforms (Non-impacted). **(B)** Proportion of protein-coding isoforms of high-risk ASD genes from Satterstrom et al., uniquely differentially expressed at isoform level, either impacted (red) or not impacted (blue) by rare *de novo* ASD LoF variants. **(C)** Ward hierarchical clustering of isoforms from panel b) based on average expression values across developmental periods. **(D)** Expression profiles of impacted and non-impacted isoforms of four ASD risk genes across development demonstrating higher prenatal expression of some impacted isoforms. **(E)** Schematic definition of alternatively regulated microexons (upper panel), proportion of all brain-expressed genes with alternatively regulated microexons (bottom left), and proportion of all brain-expressed isoforms with alternatively regulated microexons (bottom right). * - P ≤ 0.1, ** - P ≤ 0.05, *** - P ≤ 0.01.

We then selected genes with differentially expressed isoforms between adjacent development periods, for which at least one isoform is ASD LoF-impacted, and at least one other isoform is non-impacted by ASD LoF; 26 genes out of 102 Satterstrom genes satisfied this criterion (**Fig. 2B**). Hierarchical clustering of the isoforms from these genes based on expression values identified a prenatally expressed cluster consisting largely of the ASD LoF-impacted isoforms (**Fig. 2C**). A higher fraction of LoF-impacted isoforms carried microexons (i.e. short exons of 3-27bp in length) as compared to non-impacted isoforms (Permutation test, n=1,000 permutations, P = 0.04) (**Fig. 2D**), recapitulating previous findings at the gene level, and in agreement with important role of microexons in autism (Irimia et al., 2014; Li et al., 2015). The impacted and non-impacted isoforms of some genes (*KMT2C*, *MBD5*, and *PTK7)* had opposite developmental trajectories, whereas for other genes (*GABRB3*) the impacted isoforms were highly expressed throughout brain development (**Fig. 2E**). It is likely that a LoF mutation that impacts highly prenatally expressed isoform can severely disrupt early brain development and lead to NDD. In the more general case, we also found that ASD LoF-impacted isoforms were more highly expressed throughout development (P < 0.0001, Wilcoxon test), although we had not observed any specific temporal patterns (**Supplementary Figure 7).** Overall, mapping of NDD risk mutations onto the full-length isoform transcriptome could help to better understand their functional impact in the context of brain development.

### Isoform co-expression modules capture distinct trajectories of brain development

To understand how brain development is regulated at the full-length isoform level, we carried out weighted gene (and isoform) co-expression network analysis (WGCNA) (Langfelder and Horvath, 2008) (**Materials and Methods**). This analysis identified modules of genes and isoforms with highly correlated expression profiles across all BrainSpan samples. We identified a total of 8 gene and 55 isoform co-expression modules by analyzing the gene and isoform transcriptomes (**Supplementary Tables 8 and 9**). Both gene and isoform networks followed scale-free topology, and had similar low soft-thresholding beta (two for gene and three for isoform) selected to allow for module detection (**Supplementary Figure 8**).

The hierarchical clustering of modules by eigengenes demonstrated that each gene co-expression module closely clustered with a corresponding isoform co-expression module (**Fig. 3A**). Further characterization of these gene/isoform module pairs via GO annotations showed overlapping functions and pathways (**Supplementary Tables 10 and 11**). For example, gene module gM2 and isoform module iM2 were both enriched for GO terms related to synaptic transmission. This indicates that the isoform co-expression network recapitulates functional aspects of the gene co-expression network.

**Figure 3.**
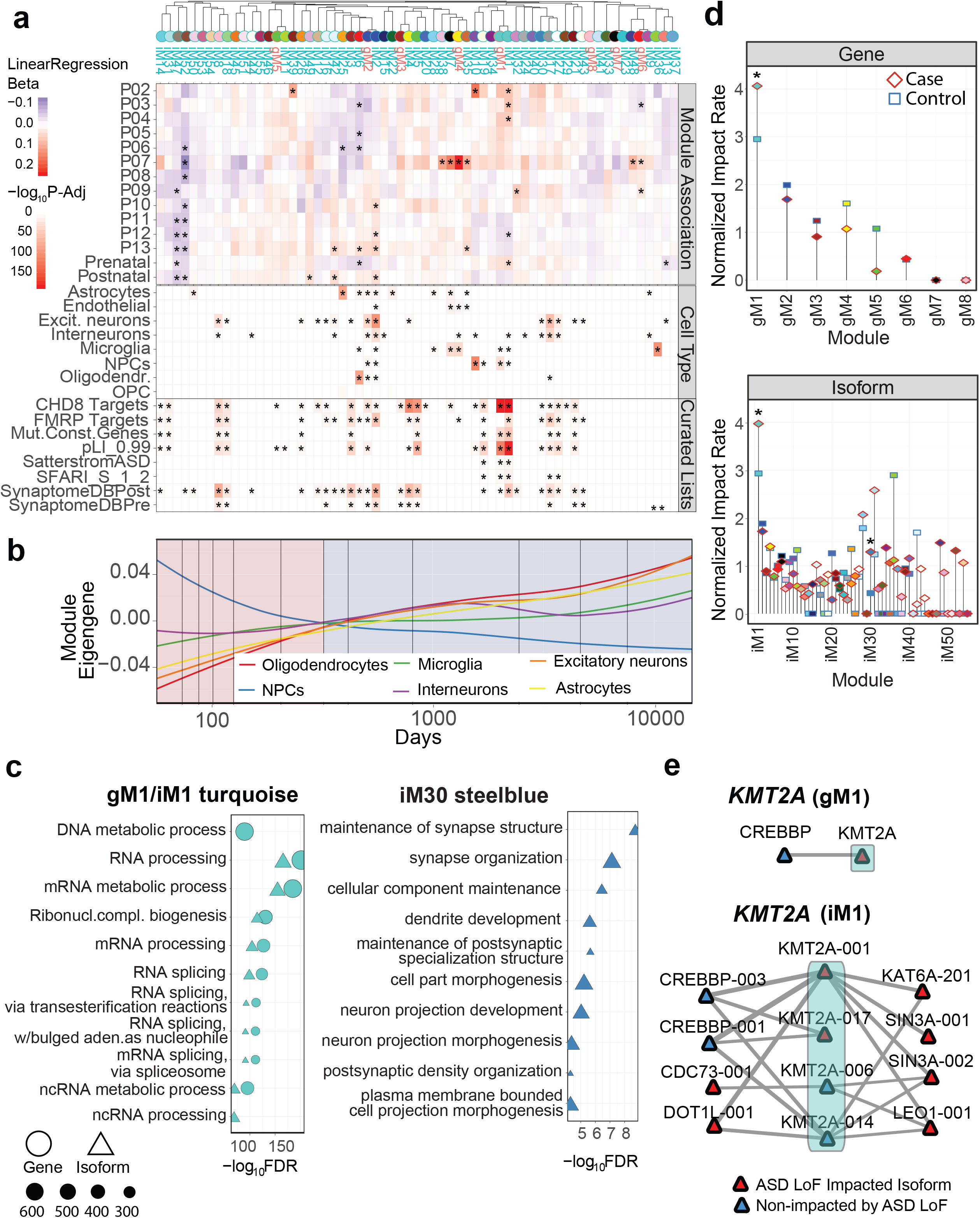
Gene and isoform co-expression analyses. **(A)** Association of gene and isoform co-expression modules clustered by module eigengene with developmental periods (top). Linear regression beta coefficients were calculated using linear mixed effect model. Module enrichment in cell type and literature curated gene sets (bottom) was calculated using Fishers exact test. **(B)** Module eigengene expression profiles across brain development for modules most significantly associated with each cell type: Astrocytes, iM25; Oligodendrocytes, iM6; Microglia, iM36; NPCs, iM10; Excitatory neurons, iM2; Interneurons, iM17. **(C)** Gene Ontology functional enrichment analyses of gM1/iM1 and iM30 modules significantly impacted by case ASD LoF mutations. **(D)** Gene (top panel) and isoform (bottom panel) co-expression modules impacted by case and control ASD LoF mutations. Normalized impact rate per module is shown. Significance was calculated by permutation test (1,000 permutations, * - FDR ≤ 0.05). **(E)** Gene-level and isoform-level co-expressed protein interaction networks for *KMT2A* gene from gM1 and iM1 turquoise modules. Only edges in the top 10% of expression Pearson correlation coefficients that are also supported by gene-level protein interactions are retained.

This hierarchical clustering of modules by eigengenes also revealed isoform modules that are completely unique relative to the gene-level modules (**Fig. 3A**). We further performed a Pearson correlation analysis between the isoform modules and the gene modules by their respective eigengene expressions and identified isoform modules with low Pearson correlation coefficients (PCC) with any gene module (**Supplementary Figure 9**). This set of isoform modules with unique expression trajectories includes those related to chemical synaptic transmission and neurogenesis (iM18, PCC=0.13); kinase activity (iM51, PCC=0.27); cell adhesion, neuron fate, and cerebral cortex regionalization (iM40, PCC=0.34); behavior, cognition, learning and memory (iM13, PCC=0.38); and chromatin, neuron organization and migration (iM31, PCC=0.3) as examples. These results identify co-expression modules with unique developmental trajectories and biological functions that were not detected by gene co-expression analyses.

In order to relate each co-expression module with brain developmental periods, we calculated module-period associations using linear mixed effects models (**Materials and Methods**). We found modules that were significantly associated with several developmental periods (**Fig. 3A, top panel**); iM1 was significantly associated with prenatal periods P02 (FDR-adjusted P = 0.009), P03 (FDR-adjusted P = 0.003), and P04 (FDR-adjusted P = 0.008; whereas iM10 and iM39 were both associated with P02 (FDR-adjusted P = 6.59E-04 and FDR-adjusted P = 0.026, respectively). Functional GO analyses of these modules demonstrated that iM1 was enriched in splicing functions, iM10 in mitosis and cell cycle-related processes, whereas iM39 was enriched in embryonic development; all functions were related to early fetal brain development (**Supplementary Table 11**). The developmental trajectory of iM10 module has a clear peak at early developmental periods, which levels down after the birth, and both modules have unique GO functions along with intermediate-to-low correlation with gene modules (**Supplementary Figure 9**).

Several other modules (gM4, iM35, iM7, and iM38) were strongly associated with the late fetal period P07 (gM4: FDR-adjusted P = 1.78E-09; iM35: FDR-adjusted P = 8.23E-04; iM7: FDR-adjusted P = 3.83E-04; iM38: FDR-adjusted P = 0.009). Collectively, these modules were enriched for angiogenesis and extracellular matrix organization GO functions (**Supplementary Table 11**), and had a high PCC and overlapping GO functions with gene modules (**Supplementary Figure 9**).

The analysis of cell-type markers extracted from single-cell sequencing studies (**Materials and Methods**) identified modules that were significantly enriched in specific cell types (**Fig. 3A, middle panel**). For example, iM10, which was associated with very early P02 period, was also enriched in neuroprogenitors (NPCs), the cells that give rise to other neuronal cell populations and often found very early in brain development. Likewise, iM2 was primarily associated with postnatal periods, and was strongly enriched in excitatory neurons, which represent mature neuronal population. Interestingly, the cluster of modules that was strongly associated with late fetal P07 period (gM4, iM35, iM7, and iM38), was enriched in microglia, or innate immune cells of the brain, that peak around late mid-fetal to late fetal development. Furthermore, isoform module eigengene trajectories captured the appropriate signals from each cell type, with NPC steadily decreasing and neuronal cell types increasing from prenatal to postnatal brain development (**Fig. 3B**). To ensure that the observed cell-type specific signatures are driven by isoform expression rather than cellular composition, we performed decomposition analyses of bulk BrainSpan RNA-seq data with fetal brain single-cell RNA-seq from two previous studies (Polioudakis et al., 2019; Zhong et al., 2018). We observed that cell type composition only explained a relatively small portion of total variance compared to sample age (i.e. developmental period) (**Supplementary Figure 10**).

Analysis of curated gene lists in the context of co-expression modules identified gM1/iM1 as being enriched in ASD risk genes, CHD8 target and functionally constrained and mutation intolerant (pLI>0.99) genes (**Fig. 3A, bottom panel)**. The same modules were significantly associated with prenatal periods and were enriched in RNA processing and splicing GO functions (**Fig. 3C, left panel).** Another module enriched in ASD risk genes was iM19 and was annotated with chromatin and histone-related GO functions. This is consistent with previous observations about enrichment of chromatin modifier genes among ASD risk genes (De Rubeis et al., 2014). In summary, the analysis of isoform co-expression modules provides novel insights and further broaden our knowledge of the developing human brain at the full-length isoform transcriptome level.

### LoF-impacted co-expression modules point to dysregulation of RNA splicing and synaptic organization

We next investigated enrichment of rare *de novo* ASD variants from cases and controls (Satterstrom et al., 2020), and identified co-expression modules that were significantly impacted by LoF case, but not control, mutations (**Supplementary Table 12**) (**Materials and Methods**). We observed three modules significantly impacted by case ASD variants: one gene module (gM1) and two isoform modules (iM1 and iM30) (**Fig. 3D**). Unsurprisingly, gM1 and iM1 clustered together and were enriched in similar GO functions related to RNA processing and splicing, including non-coding RNA splicing (**Fig. 3C**). This agrees with the already-demonstrated crucial role of splicing dysregulation in ASD (Gandal et al., 2018; Parikshak et al., 2016). Functional enrichment of isoform co-expression module iM30 pointed to dysregulation of synapse organization, dendrite development and neuronal projection pathways (**Fig. 3C**), which are strongly implicated in ASD (Iakoucheva et al., 2019; Pinto et al., 2014). The highest PCC of iM30 with any gene module is 0.62, and its developmental trajectory and GO functions are distinct when compared to gene modules **(Supplementary Figure 9)**. Thus, isoform modules reflect processes previously implicated in ASD, and point to specific isoforms (rather than genes) that can contribute to this dysregulation.

To demonstrate how isoform co-expression modules could be useful for future studies, we built isoform co-expressed protein-protein interaction (PPI) networks for the gM1 and iM1 modules (**Supplementary Figure 11**). The isoform network was focused on ASD risk genes that had at least one isoform impacted by an LoF mutation, and the edges that had gene-level PPI information (due to scarcity of isoform-level PPIs) were filtered for the top 10% PCC (**Materials and Methods**). Clearly, gM1 had fewer connections than iM1, and iM1 highlighted some interesting isoform co-expressed PPIs that were not detectable from the gene-level network. For example, 9 genes from this module (*ARID1B*, *CHD8*, *KMD5B*, *KMT2A*, *MED13L*, *PCM1*, *PHF12*, *POGZ*, and *TCF4*) had at least one ASD LoF-impacted isoform and at least one that was not impacted by mutation. These isoforms were co-expressed and interacted with different partner isoforms. For example, ASD LoF-impacted and non-impacted isoforms of *KMT2A* gene had shared as well as unique protein interacting partners (**Fig. 3E**). This could lead to different networks being disrupted as a result of ASD mutation, and these networks were not observed at the gene level, with only one *KMT2A* partner (*CREBBP*) in the gene network. Another interesting observation from co-expressed PPI networks was that LoF-impacted isoforms tended to have higher correlation with corresponding partners than non-impacted isoforms (Mann-Whitney test, P= 1.53E-05), suggesting potentially greater functional impact on networks.

### *De novo* splice site mutations of NDD risk genes cause exon skipping, partial intron retention, or have no effect on isoforms

One type of LoF mutation is mutations that affect splice sites directly. Here, we experimentally investigated the effect of *de novo* splice site mutations identified by exome sequencing studies in four NDD risk genes (*DYRK1A, SCN2A, DLG2,* and *CELF2)* to better understand their functional impact. All highly prenatally expressed isoforms of these genes were found in iM1. We used an exon trapping assay (**Materials and Methods**) to test the following *de novo* splice site mutations: *SCN2A* (chr2:166187838, A:G, acceptor site) (Fromer et al., 2014); *DYRK1A* (chr21: 38865466, G:A, donor site) (O’Roak et al., 2012); *DLG2* (chr11: 83194295, G:A, donor site) (Fromer et al., 2014); and *CELF2* (chr10: 11356223, T:C, donor site) (Xu et al., 2011). The mutation in *SCN2A* caused out-of-frame exon skipping and potential inclusion of 30 new amino acids into the translated protein before ending with a premature stop codon that most likely will result in nonsense-mediated decay (NMD) (**Fig. 4A**). In contrast, the mutation in *DYRK1A* caused an in-frame exon skipping, potentially producing a different variant of the same protein and thus is expected to have milder or perhaps no functional effect (**Fig. 4B**). In the case of *DLG2*, the mutation affected a splice site adjacent to the exon five, which is alternatively spliced in the WT isoforms (**Fig. 4C**). We constructed a minigene that includes exon five together with the preceding exon four and observed that exon five was constitutively spliced out from our construct independently on the presence of mutation. However, the mutation caused partial (i.e., 65bp) intron inclusion downstream from exon four. At the translational level, this mutation would likely result in a truncated protein one residue after the end of exon 4 due to a premature stop codon. Finally, the *CELF2* mutation affected an alternative splice site, which also mapped to an exonic region of another alternatively spliced isoform. When cloned into the exon trapping vector, the isoform generated from the WT minigene included the isoform carrying longer exon with mutation (**Fig. 4D**). Thus, after introducing the mutation, no difference between WT and mutant constructs was observed. This is not surprising given the fact that the splice site mutation behaves like exonic missense mutation in the isoform predominantly expressed from our construct. These results suggest that mutations could impact different isoforms of the same gene by different mechanisms, i.e., splice site mutation in one isoform could represent a missense mutation in another isoform.

**Figure 4.**
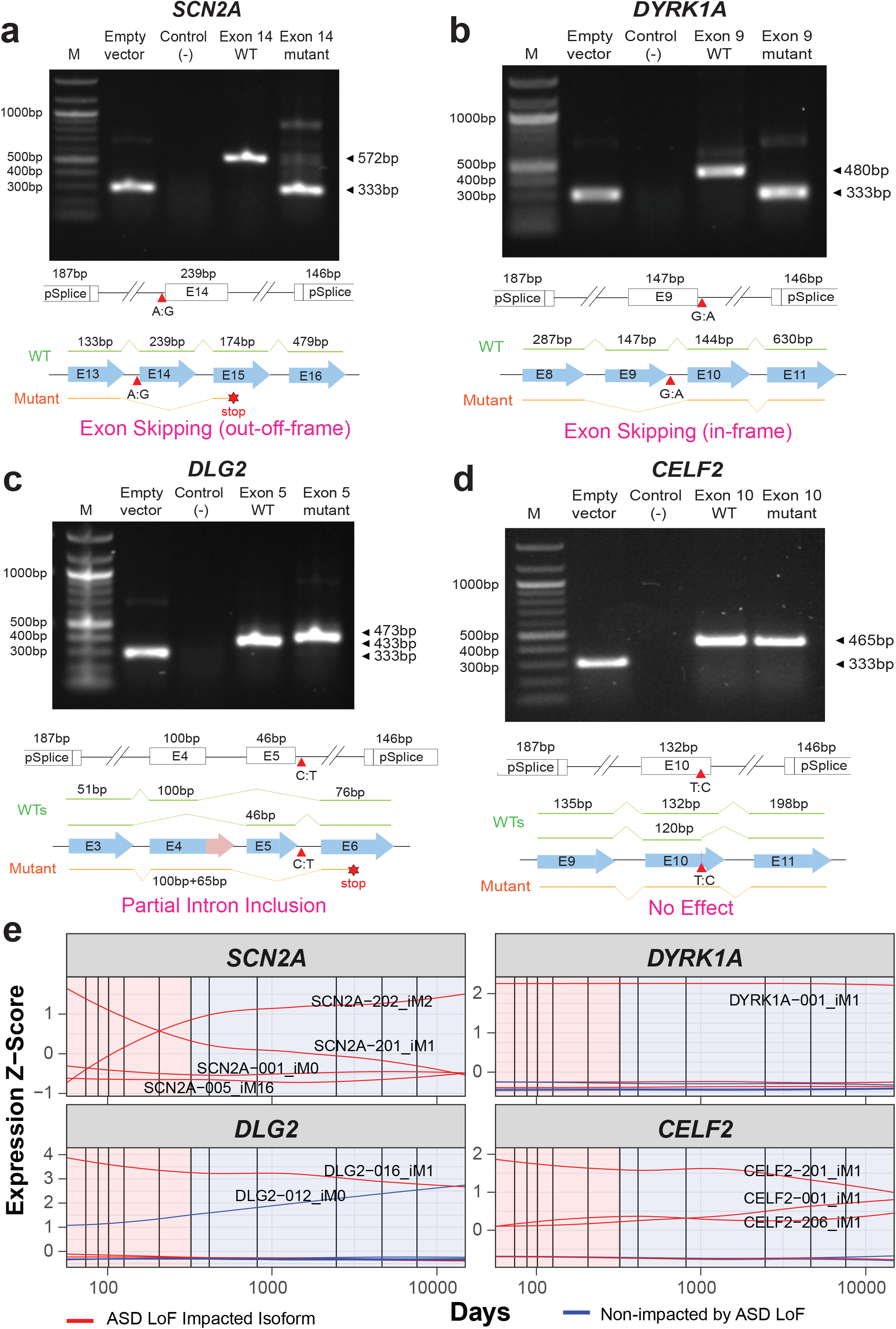
Functional effect of the de novo splice site mutations from the patients with neurodevelopmental diseases. Minigene assays demonstrate the effect of splice site mutations in four genes. **(A)** *SCN2A*; **(B)** *DYRK1A*; **(C)** *DLG2*; and **(D)** *CELF2*. Schematic representation of the cloned minigenes, the expected splicing patterns, and the impact of the mutations are shown below the gel image. Numbers denote base pairs; M: molecular marker; E: exon. **(E)** Expression profiles across brain development of the brain-expressed isoforms transcribed by these four genes, annotated with module memberships; highly overlapping expression profiles are unlabeled for readability.

Further analysis of expression profiles of the brain-expressed isoforms transcribed by these genes (**Fig. 4E**) suggested that highly prenatally expressed isoforms (*SCN2A-201, DYRK1A-001, DLG2-016* and *CELF2-201*) were most likely targets for the “pathogenic” effect of mutations. Furthermore, given distinct co-expressed PPI partners of impacted *vs* non-impacted isoforms (**Supplementary Figure 12**) in most cases, the effect of mutation would be propagated onto different networks, affecting different signaling pathways. In summary, our experiments showcased different scenarios of the impact of splice site mutations and confirmed the need to investigate their functional impact at the isoform-rather than the gene-level resolution.

### The splice site mutation in *BTRC* reduces its translational efficiency

As shown above, mutations can impact different protein interaction networks depending on the splicing isoform that they affect, and whether the impacted isoform is expressed at specific periods of brain development. Next, we investigated in greater detail how specific mutations mapped to different isoforms may disrupt downstream signaling pathways. For this, we selected three full-length isoforms (*BTRC-001, BTRC-002,* and *BTRC-003*) of an ASD risk gene, *BTRC* (also known as *β-TrCP* or *FBXW1A*) (Ruzzo et al., 2019), based on their availability from our previous study (Yang et al., 2016) (**Fig. 5A**). Two *de novo* mutations, one missense (chr10:103285935,G-A)(Ruzzo et al., 2019) and one splice site (chr10: 103221816, G:A, donor site)(De Rubeis et al., 2014), were identified in ASD patients with zero in controls, making *BTRC* one of 69 high-confidence ASD risk genes with genome-wide significance 0.05 < FDR ≤ 0.1 (Ruzzo et al., 2019). We demonstrated that the splice site *BTRC* mutation caused in-frame exon four (78bp) skipping in the exon trapping assay (**Fig. 5B**). To further test the effect of this mutation on different *BTRC* transcripts, we generated additional constructs by inserting abridged introns surrounding exon 4 into the coding sequence (CDS) of two isoforms, *BTRC-001* and *BTRC-002* (**Fig. 5C**, **Materials and Methods**). The third isoform, *BTRC-003*, did not carry exon 4, and its structure and size are identical to the *BTRC-001*, after exon 4 is skipped. We also generated mutant constructs BTRC-001Mut and BTRC-002Mut carrying the mutation in the abridged intron (**Fig. 5A**). The RT-PCR following exon trapping assays on the full-length CDS, as well as WT and mutant constructs with abridged introns, confirmed the correct sizes of all constructs, and validated exon skipping event due to splice site mutation (**Fig. 5C**). The expression levels of the mutant constructs were comparable to the WT. Furthermore, Western blot confirmed the expected sizes of the protein products produced from the WT and mutant constructs (**Fig. 5D**). The splice site mutation significantly reduced the amount of the protein produced from mutant isoforms, suggesting their decreased translational efficiency (**Fig. 5E**). Higher amount of protein product produced from all constructs with abridged introns compared to CDSs was consistent with previous observations of increased translational efficiency of RNAs produced by splicing compared to their intron-less counterparts (Diem et al., 2007). Further, *BTRC-001* and *BTRC-002* are highly expressed relative to the non-impacted *BTRC-003* (**Fig. 5F**) and are co-expressed and interact with non-overlapping sets of protein partners in the co-expressed PPI networks (**Supplementary Figure 13**). This suggests that mutations in different isoforms could impact different cellular networks.

**Fig. 5.**
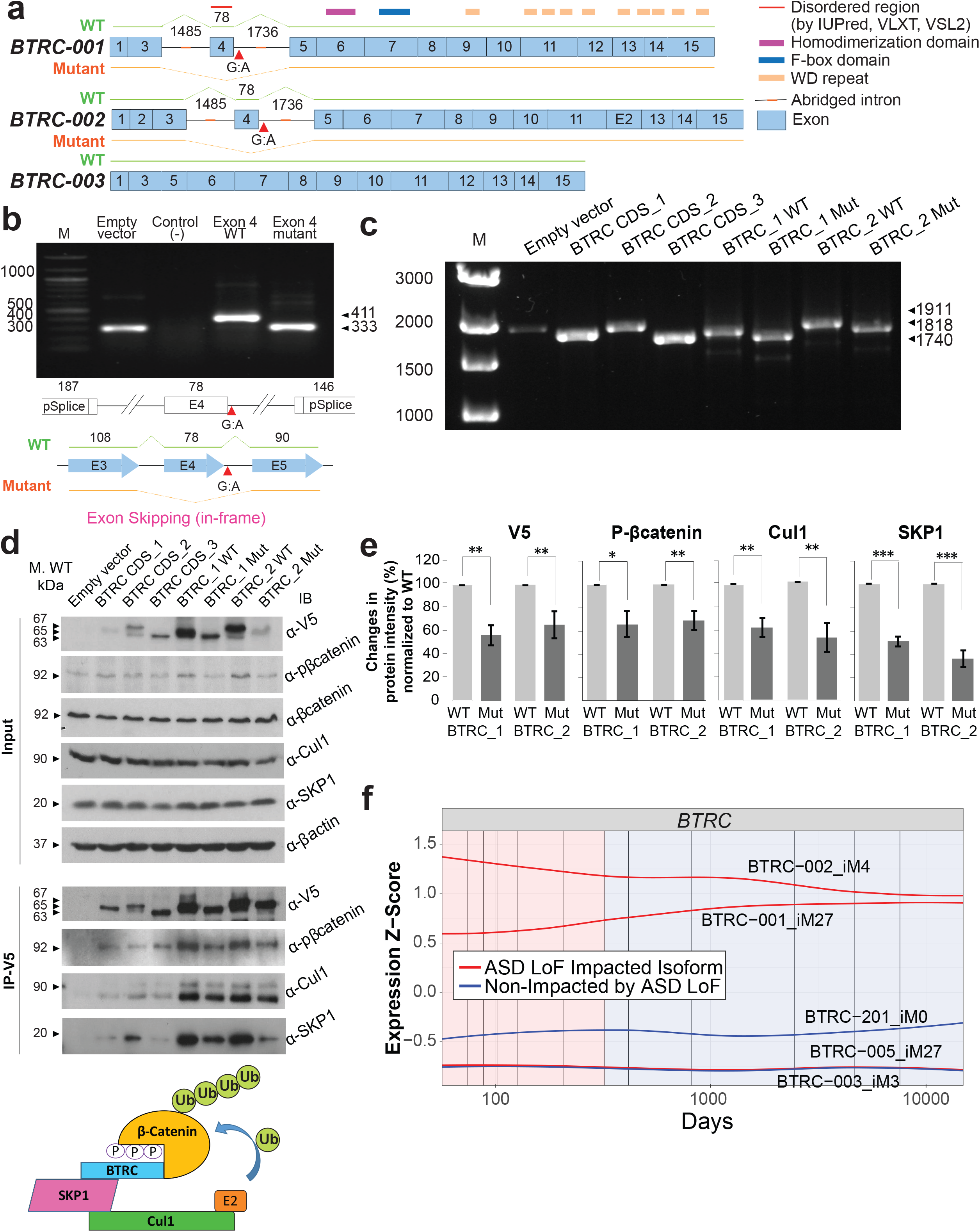
The *de novo* autism splice site mutation causes exon skipping in *BTRC* isoforms and reduces their translational efficiency. **(A)** The exon structure of three splicing isoforms of the *BTRC* gene showing positions of the cloned abridged introns and the splice site mutation; numbers denote base pairs (bp). **(B)** Minigene assays demonstrate exon 4 skipping as a result of the splice site mutation. The assays show the RT-PCR results performed using total RNA from HeLa cells transfected with *BTRC* minigene constructs; numbers denote base pairs. **(C)** Splicing assays with the full-length constructs carrying abridged introns confirm exon skipping observed in the minigene assays. **(D)** Immunoblotting (IB) from the whole cell lysates of HeLa cells transfected with different BTRC minigene constructs and an empty vector, as indicated. Membranes were probed to observe BTRC overexpression, and to investigate expression of p-β-catenin, Cul1 and SKP1. β-actin was used as loading control. Immunoprecipitation was performed with the antibody recognizing V5-tag and proteins were detected by immunoblotting (IB) with the p-β-catenin, Cul1, SKP1 and V5 antibodies. The splice site mutation causes reduced translational efficiency of both BTRC_1Mut and BTRC_2Mut mutant isoforms as compared to their wild type counterparts. Schematic diagram of *Skp1-Cul1-BTRC* ubiquitin protein ligase complex is shown at the bottom. **(E)** Quantification of protein pull-downs with V5-IP using ImageJ software. The band intensity values were normalized to WT expression levels. Error bars represent 95% confidence intervals (CI) based on 3 independent experiments. On average, 40% reduction of *BTRC* protein expression is observed as a result of a mutation. Consequently, the reduction of the corresponding BTRC binding partners (p-β-catenin, Cul1, and SKP1) is also observed. **(F)** Expression profiles of brain-expressed *BTRC* isoforms show higher expression of ASD-impacted BTRC-001 and BTRC-002. Numbers denote base pairs (a, b, c panels) or kDa (d). P-values: * - P<0.05, ** - P ≤ 0.01, *** - P ≤ 0.001.

Next, we investigated binding properties of all isoforms using co-immunoprecipitation (co-IP) (**Fig. 5D**). The *BTRC* gene encodes a protein of the F-box family and is a component of the SCF (Skp1-Cul1-F-box protein) E3 ubiquitin-protein ligase complex. One of the well-known substrates of this complex is β-catenin (CTNNB1). SCF complex ubiquitinates and regulates degradation of β-catenin, an essential component of the *Wnt* signaling pathway (Winston et al., 1999). *Wnt* plays key roles in cell patterning, proliferation, polarity and differentiation during the embryonic development of the nervous system (Ciani and Salinas, 2005), and it has been consistently implicated in ASD (Iakoucheva et al., 2019; Kwan et al., 2016). Both β-catenin and Cul1 carry *de novo* mutations identified in patients with NDD (Satterstrom et al., 2020).

The interaction of BTRC with its partners, Cul1, Skp1 and β-catenin, demonstrated reduced binding with mutant BTRC, potentially suggesting shortage of the SCF ligase complexes (**Fig. 5D-E**). In agreement with previous observations, we found that BTRC only binds to the phosphorylated form of β-catenin (Winston et al., 1999). This suggests that the amount of protein complex is strongly dependent on the availability of *BTRC* protein, which is significantly reduced due to splice site mutation. Thus, our results indicate that the *BTRC* splice site mutation causes exon skipping in *BTRC* isoforms and reduces translational efficiency of the resulting protein product. This, in turn, decreases the amount of SCF protein ligase complexes that are available for β-catenin ubiquitination. We hypothesize that this may lead to impaired degradation of β-catenin, its cellular accumulation and upregulation of *Wnt* signaling as a result of this ASD risk mutation. Further studies in neuronal cells are needed to test this hypothesis.

## Discussion

Recent large-scale whole exome and whole genome sequencing studies have greatly facilitated the discovery of the genetic causes of neurodevelopmental disorders. One of the bottlenecks in translating these findings into molecular mechanisms is our limited understanding of the transcriptional and translational programs governing brain development. The brain is one of the most complex human organs with the highest number of alternatively spliced events (Mele et al., 2015; Raj and Blencowe, 2015). Thus, the knowledge of its splicing repertoire is crucial for future translational studies in brain diseases.

Several earlier studies had addressed the impact of NDD mutations at the exon-level resolution. For example, Uddin *et al*. identified highly expressed critical exons with *de novo* ASD mutations that were enriched in cases compared to controls (Uddin et al., 2014). A more recent study correlated exons impacted by ASD LoF mutations with patients’ phenotypes, and found that patients with mutations in the same exon had more similar phenotypes (Chiang et al., 2020). However, none of previous studies investigated the impact of LoF mutations in the context of the full-length isoform transcriptome.

We previously demonstrated that integration of genetic data with isoform-level co-expression and protein interaction networks were crucial for improving understanding of the molecular mechanisms of neurodevelopmental disorders (Corominas et al., 2014; Gandal et al., 2018; Lin et al., 2015; Lin et al., 2017). The importance of isoform-level networks was further emphasized by the observation that protein products encoded by different splicing isoforms of the same gene share less than half of their interacting partners (Yang et al., 2016). These studies underscore the importance of the brain isoform transcriptome for future studies of neurodevelopmental diseases. Here, we analyzed the full-length isoform transcriptome of the developing brain and demonstrated its utility for investigating loss-of-function mutations implicated in autism.

We demonstrated that brain differential isoform expression analysis identified a fairly large set of DEIs that were not detected by the gene-level analysis. Furthermore, DEIs captured more relevant functions than DEGs in the context of brain development. Processes such as neuron projection development, axon development, head, brain and nervous system development were primarily supported by DEIs uniquely identified only through analysis of the isoform transcriptome.

By mapping LoF mutations from autism cases and controls onto the full-length isoform transcriptome, we found that ASD LoF-impacted isoforms had significantly higher prenatal expression than non-impacted isoforms or isoforms impacted by control mutations. The expression trajectories of impacted and non-impacted isoforms across brain development were remarkably different for some of the autism risk genes. For example, two LoF-impacted isoforms of *KMT2C* histone lysine methyltransferase, a high-confidence ASD risk gene (De Rubeis et al., 2014; Iossifov et al., 2015; O’Roak et al., 2011) were highly expressed prenatally and had opposing temporal trajectories compared with non-impacted isoforms that were highly expressed postnatally (**Fig. 2D**). Similar pictures were observed for *PTK7* and *MBD5*. This demonstrates that future studies of these and other genes with similar properties should focus on impacted isoforms with high prenatal expression, rather than on all available isoforms, as they may be more relevant to brain development.

In general, we consistently gain additional information from the full-length isoform transcriptome across various types of analyses. At the level of co-expression, isoform co-expression modules provide important insights into neurodevelopment and on how it may be disrupted by autism mutations. Importantly, many isoform modules, especially those with low PCC with gene modules, have unique and distinct developmental trajectories and biological functions that are not identified through gene level analyses (**Supplementary Figure 9**). Many of these distinct isoform modules were annotated with important and ASD-relevant functions, such as translation (iM21), protein/histone demethylation (iM47), chromatin organization and neuron migration (iM31), and cell adhesion, neuron fate, cerebral cortex regionalization (iM40). Other isoform modules that have high PCC with gene modules, could also provide important insights. The isoform module iM1 was significantly enriched in isoforms impacted by case LoF mutations (**Fig. 3D**). Functionally, it was enriched in RNA splicing and processing pathways that have been previously implicated in ASD (Parikshak et al., 2016). iM1 was significantly associated with prenatal developmental periods; enriched in interneurons, microglia and NPCs; enriched in CHD8 target genes, mutation intolerant genes, and was also one of a few modules enriched in ASD risk genes. Given all these lines of evidence, it clearly is a very important module for further investigation.

Using isoforms from the iM1 module, we experimentally investigated the functional impact of splice site-disrupting LoF mutations in five genes. The results demonstrated exon skipping or disruption of normal splicing patterns, albeit not in all cases. A more detailed analysis at the isoform level suggested that not all isoforms were affected by mutations. For example, at least one known isoform of the *BTRC* gene did not carry an exon with mutation, and therefore was not expected to be impacted by it. We next demonstrated that a *BTRC* mutation decreased translational efficiency of the impacted isoforms, since lower amount of the resulting protein was observed (**Fig. 5D**). This, in turn, lead to reduced interaction between BTRC and its protein partners, potentially disrupting *Wnt* signaling **(Fig. 5E**). Since β-catenin is a substrate of the *BTRC-Cul1-Skp1* ubiquitin ligase complex, the shortage of this complex may lead to impaired ubiquitination and degradation of β-catenin and its neuronal accumulation. Interestingly, transgenic mice overexpressing β-catenin have enlarged forebrains, arrest of neuronal migration and dramatic disorganization of the layering of the cerebral cortex (Chenn and Walsh, 2002). It would be interesting to investigate whether the patient carrying the *de novo BTRC* splice site mutation has similar brain abnormalities.

Typically, mutations affecting essential splice sites are automatically classified as LoF mutations when considering gene-level analyses. Here, we demonstrate that this is not always the case, and that splice site mutations impacting one isoform of a gene may serve as a missense mutation in another isoform that carries a longer exon spanning the splice site, like in the case of *CELF2* (**Fig. 4D**). Thus, depending on where, when, at what level and which isoform of the gene is expressed, the functional impact of the same mutation may differ dramatically. In addition, the mutation could also be “silent” if the isoform is highly expressed but does not carry an exon affected by a specific mutation. This suggests that the impact of mutations should be investigated at the isoform-rather than the gene-level resolution, and expression levels of splicing isoforms in disease-relevant tissues should be taken into consideration to better guide hypotheses regarding potential mechanisms of the disease and its future treatments.

While our study provided new insights into functional impact of NDD mutations at the level of the full length isoform transcriptome, it has several limitations that warrant a discussion. The BrainSpan developmental brain transcriptome was sequenced using short read sequencing technologies, and therefore full-length isoform assignment, especially for low-abundant transcripts, is less reliable as would be expected from the long read sequencing technologies. Long read RNAseq could offer large gains in the accuracy of full-length isoform identification and discovery. When it becomes less cost-prohibitive, creating a long read transcriptome of the developing human brain resource would be invaluable for the scientific community. In summary, our study directly demonstrates the value of using full length transcriptome data for future human disease studies and for investigating functional impact of disease-associated mutations.

### Data and code availability

Raw RNAseq isoform-level BrainSpan data are available at PsychENCODE Capstone Data Collection, www.doi.org/10.7303/syn12080241. The processed summary-level BrainSpan data are available at http://Resource.PsychENCODE.org. The code used for isoform RNAseq data analysis generated during this study is available from GitHub (https://github.com/IakouchevaLab/Isoform_BrainSpan).

## Acknowledgements

The study was supported by the grants R01MH109885 (LMI), R01MH105524 (LMI), R01 MH108528 (LMI) and, in part, by the Simons Foundation for Autism Research (345469, LMI). We thank Dr. Marc Vidal and Dr. David Hill for providing the clones of *BTRC* isoforms. We also thank Dr. Hyun-Jun Nam and Bom Han from Iakoucheva lab for helping with genetic data curation.

## Author contributions

L.M.I., G.N.L., and R.C. conceived and conceptualized the study; K.C., P.Z., A.B.P., and J. C. performed bioinformatics analysis; J.U., M.A., and A.T. performed experiments; L.M.I. supervised the work and acquired funding; K.C and L.M.I. wrote the paper with input from all authors.

## Materials and Methods

R version 3.6.0 was used throughout this analysis. Downstream bioinformatics analysis is outlined in **Supplementary Figure 1A.**

### Processing of RNA-Seq gene and transcript BrainSpan data

RNA-Seq datasets quantified at the gene and isoform levels were downloaded from PsychENCODE Knowledge Portal, PEC Capstone Collection, Synapse ID: syn8466658 (https://www.synapse.org/#!Synapse:syn12080241). RNA-Seq from post-mortem brain tissue of 57 donors aged between 8 weeks post-conception to 40 years, across a number of different brain regions, for a total of 606 samples, has been carried out as previously described (Li et al., 2018) (**Supplementary Figure 1B**). Data processing was performed as described (Gandal et al., 2018). Briefly, FASTQs were trimmed for adapter sequence and low base call quality (Phred score < 30 at ends) using cutadapt (v1.12). Trimmed reads were then aligned to the GRCH37.p13 (hg19) reference genome via STAR (2.4.2a) using comprehensive gene annotations from Gencode (v19). BAM files were produced in both genomic and transcriptome coordinates and sorted using samtools (v1.3). Gene and isoform-level quantifications were calculated using RSEM (v1.2.29). Quality control metrics were calculated using RNA-SeQC (v1.1.8), featureCounts (v1.5.1), PicardTools (v1.128), and Samtools (v1.3.1). Subsequently, TPM matrices for both, gene and transcript datasets, were filtered for TPM ≥ 0.1 in at least 25% of samples, yielding a total of 100,754 isoforms corresponding to 26,307 genes.

Sample connectivity analysis was performed to detect sample outliers as previously described (Oldham et al., 2012). In brief, bi-weight mid-correlation was calculated among sample expression vectors in both filtered datasets. These values were converted into connectivity Z-scores. 55 samples were identified as having sample connectivity Z-scores ≤ -2, and were removed from downstream analysis, resulting in 551 final samples.

Surrogate variable analysis (SVA) was performed to remove latent batch effects in the data, taking into consideration age, brain region, sex, ethnicity, and study site (Leek, 2014; Leek and Storey, 2007). The number of surrogate variables was chosen to minimize apparent batch effects while avoiding overfitting based on evidence from principal components analysis and relative log expression (**Supplementary Figures 2-5**). 16 surrogate variables were found to be sufficient for downstream analysis of both gene and transcript data.

### Validation of isoform expression with RT-qPCR

RT-qPCR was used to estimate the relative expression difference of 44 unique isoforms of 32 genes between two independent RNA samples that were age, sex and brain region-matched to the samples from the BrainSpan; these results were then compared to the computationally-assigned BrainSpan values (**Supplementary Table 1**, **Supplementary Figure 6**). 24 out of 44 isoforms were derived from 12 genes (i.e. two isoforms per gene). RNA from a frontal lobe tissue sample of a 22 weeks old male (fetal brain), and RNA from cerebral cortex tissue sample of a 27 years old male (adult brain) (AMSBIO, UK), corresponding to P06 (late mid-fetal) and P13 (young adult) in the BrainSpan data, was used. The BrainSpan isoform expression data was then compared to the RT-qPCR experimental expression results. The isoforms were selected to encompass all possible ranges of differential expression - from high differential expression (high log2FC, *e.g.* > 4), to low or no differential expression (low log2FC, *e.g.* < 1) between the same isoforms from two samples from different time points. To further discriminate between isoform expression and absolute gene expression, isoforms were selected based on the presence of unique exons, for which the primers were designed. The majority of selected isoforms were expressed with >1 TPM values to ensure that they were detectable by the RT-qPCR. However, we also included 12 isoform pairs with at least one isoform being lowly expressed (<1 TPM) to ensure that validation covers both highly and lowly expressed isoforms. This resulted in a total of 44 unique isoforms from 32 genes. Primers were designed using unique exon-exon junctions, specific for each of the selected isoforms. 3 µg of RNA using SuperScript II Kit (Invitrogen) were reverse transcribed to cDNA, following manufacturer’s instructions. Then, the cDNA was diluted ten times to use as a template for the RT-qPCR reaction. SYBR Green II Master Mix (Invitrogen) was used for the RT-qPCR reaction, performed in a CFX Connect 96X Thermal Cycler, using standard parameters for SYBR Green. Relative expression between each isoform in the two samples was calculated by normalizing each expression value against two housekeeping genes (*RPL28* and *MRSP36*) as control using QIAGEN control primers, and ΔΔt method was applied using the CFX Manager Software. Comparison of these relative expression values against the BrainSpan computational expression values resulted in positive correlation (**Supplementary Figure 6**).

### Differential gene and isoform expression analysis

Differential gene and isoform expression analysis was performed using the *limma* (v3.40.6) R package (Ritchie et al., 2015). Relevant covariates (brain region, sex, ethnicity, and study site) and surrogate variables were included in the linear model as fixed effects. The *duplicateCorrelation* function was used to fit the donor identifier as a random effect to account for the nested expression measurements due to multiple brain regions derived from the same donor. Genes and isoforms with an absolute fold change of ≥1.5 and FDR-adjusted p-value of ≤0.05 between adjacent developmental periods, or between prenatal and postnatal periods (PrePost) were defined as significantly differentially expressed.

### Cell type and literature curated gene sets enrichment analyses

Fisher-exact tests were performed on gene lists and isoform lists (converted to gene identifiers) against curated gene lists: mutational constraint genes (Mut. Const. Genes) (Samocha et al., 2014), FMRP target genes (Darnell et al., 2011), high-risk ASD genes (Satterstrom ASD) (Satterstrom et al., 2020), CHD8 target genes (Wilkinson et al., 2015), synaptic genes (Synaptome DB) (Pirooznia et al., 2012), genes intolerant to mutations (Pli_0.99) (Lek et al., 2016), syndromic and rank 1 and 2 ASD risk genes (SFARI_S_1_2) (https://gene.sfari.org/). Cell types were extracted from a recent gene-level single cell sequencing study (Zhong et al., 2018).

### Gene Ontology (GO) functional enrichment analyses

Functional enrichment analysis was performed using the *gprofiler2* v0.1.5 R package. Ensembl gene or isoform (converted to gene) identifiers were used to test for enrichment in two Gene Ontology categories, Biological Processes (BP) and Molecular Functions (MF). Enrichment p-values were Benjamini-Hochberg corrected for multiple hypothesis testing, and overly general terms (i.e., terms with more than 1,000 members) were filtered out.

### Rare *de novo* ASD loss-of-function variants

Rare *de novo* variant data was downloaded from Satterstrom et al. (Satterstrom et al., 2020), and was processed using Ensembl’s Variant Effect Predictor v96 (McLaren et al., 2016) using human genome version GRCh37 to annotate variants for predicted functional consequences. Loss-of-function (LoF) variants were defined as those impacting essential splice donor/acceptor sites, frameshift insertions or deletions, predicted start losses, and predicted stop gains.

### Weighted gene/isoform co-expression network analyses (WGCNA)

Co-expression networks were constructed using the *WGCNA* (v1.68) R package (Langfelder and Horvath, 2008). Relevant covariates (brain region, sex, ethnicity, and study site) and surrogate variables were first regressed out of both gene and isoform expression datasets using linear mixed effects models. Each transformed expression matrix was then tested for scale-free topology to estimate a soft-thresholding power, *beta*. We used *beta=*2 for gene co-expression and *beta*=3 for isoform co-expression networks, and signed networks were constructed blockwise using a single block for the gene network and three blocks for the isoform network with deepSplit=2 and minModuleSize=20 for module detection in both networks. Both networks followed scale-free topology (**Supplementary Figure 8**).

### Co-expression module characterization

Module eigengene and developmental period association analysis was performed using linear mixed effects models, considering fixed effects (age, brain region, sex, ethnicity, and study site) and random donor effects to account for multiple brain region samples per donor. Module enrichment analysis was performed using Fisher’s exact tests against curated gene lists; isoform identifiers within modules were converted to gene identifiers for this purpose. Gene Ontology functional enrichment analysis for modules was performed using *gprofiler2* with queries ordered by module membership (kME) rank.

### Decomposition of bulk BrainSpan data with fetal brain single cell RNA-seq

Bisque (Jew et al., 2020) was used to decompose bulk RNA-seq BrainSpan data to estimate the cell type proportions. Single-cell RNA-seq data of human fetal prefrontal cortex from two studies (Polioudakis et al., 2019; Zhong et al., 2018) was used to compute the reference profile. BrainSpan data was compared to the reference profile to estimate the cell type proportion in each sample. Variance partition (Hoffman and Schadt, 2016) was used to investigate cell type proportions (**Supplementary Figure 10**).

### Variant impact analysis for co-expression modules

We quantified the impact of rare *de novo* LoF case and control variants on co-expression modules by mapping variants to the members of each module. To account for gene and isoform sizes in each module, we calculated the non-overlapping length, in base pairs, of the members of each module and normalized the module impact rates by these genomic coverages. Further, to control for total mutation rate, we scaled the impact rate by the total number of variants. We further added a scaling factor of 1,000,000 to make the numbers more manageable, in a similar fashion to a TPM calculation. Differences in the impact rates between case and control variants for each module were tested using permutation test with 1,000 iterations of module member resampling (controlling for length and GC content, ±10% for each attribute) (**Supplementary Table 12**). Modules impacted by significantly more case mutations were identified.

### Integration of protein-protein interaction networks with co-expression modules

Gene-level PPI network data was manually curated and filtered for physical and co-complex interactions extracted from Bioplex (Huttlin et al., 2015), HPRD (Keshava Prasad et al., 2009), Inweb (Li et al., 2017), HINT (Das and Yu, 2012), BioGRID (Chatr-Aryamontri et al., 2017), GeneMANIA (Zuberi et al., 2013), STRING (Szklarczyk et al., 2017), and CORUM (Ruepp et al., 2010). To build co-expressed PPI networks, gene and isoform modules were first filtered for connections (i.e. edges) supported by the gene-level PPIs; isoform edges were retained if corresponding gene edges were supported by PPIs. Subsequently, networks were filtered to only retain edges supported by the top 10% of co-expression PCCs between genes or isoforms. Genes or isoforms without any connections were removed from the networks.

### Minigenes cloning

The following genes impacted by rare *de novo* splice site mutations identified in NDDs patients were selected for the experiments: *SNC2A* (chr2:166187838, A:G, acceptor site) (Fromer et al., 2014); *DYRK1A* (chr21: 38865466, G:A, donor site) (O’Roak et al., 2012), *CELF2* (chr10: 11356223, T:C, donor site) (Xu et al., 2011), *DLG2* (chr11: 83194295, G:A, donor site) (Fromer et al., 2014) and *BTRC* (chr10: 103221816, G:A, donor site) (De Rubeis et al., 2014). The exons of these genes that are likely impacted by splice site mutations, together with the ∼1kb of their flanking intronic sequence, were cloned. The constructs were cloned into pDESTSplice exon trapping expression vector (Kishore et al., 2008). The site-directed mutagenesis by two-step stich PCR was performed to introduce the mutation affecting the splice site.

The minigenes were generated by PCR-amplifying the desired sequences from genomic DNA (Clontech). Primers were designed for each minigene, and attB sites were added at the 5’ end of the primers. The sequences of the primers were as follows: (1) *SCN2A*; Fw: GGAAGCTATGTTTAGCCAGGATACATTTGG, Rv: CCAGATGATGTCCCCTCCCTACATAGTCC; (2) *DYRK1A*: Fw: GTTGGGAAAATTTCCCCCTATTTAAGC, Rv: CCCAGAGGCTTAATAAAGTATGGACC; (3) *CELF2*: Fw: GGAGTTGGAATGACAGACGTTCACATGC, Rv: CCGCTGTGGGCTGAGGATCAGTTTCC; (4) *DLG2*: Fw: GAGGTTCAGAGACATTCAATTCCC, Rv: CTTGATGCTGTCCAGATAATGC; (5) *BTRC*: Fw: GGGCCTCAGAATGACACAGTACG, Rv: GAACTTGCGTTTCTTGTTTTTGCC. After PCR amplification, amplicons were loaded in a 1% low EEO agarose gel (G-BioSciences) and purified using the QIAquick Gel Extraction Kit (QIAGEN) following manufacturer’s instructions. Purified amplicons were subcloned into pDON223.1 expression vector using the BP-Gateway System (Invitrogen). At least six different clones for each minigene were sequenced to verify correct sequences of the minigenes. The clone with the desired sequence and highest DNA concentration was used for subcloning into the pDESTSplice expression vector (Addgene) using the LR-Gateway System (Invitrogen).

### Exon trapping and RT-PCR

HeLa cells were seeded at 2x10^5^ cells per well in 6-well plates (Falcon). After 24h, cells were transfected using Lypofectamine 3000 (Invitrogen) following manufacturer’s instructions, and then harvested after additional 24h. RNA was purified using the RNAeasy Mini Kit (QIAGEN) following manufacturer’s instructions. Two µg of RNA was used to generate cDNA using the SuperScript III First Strand kit (Invitrogen), and PCR was carried out. In the case of exon trapping assays, we used primers specific for the rat insulin exons constitutively present in the pDESTSplice vector: Fw: CCTGCTGGCCCTGCTCA, Rv: TAGTTGCAGTAGTTCTCCAGTTGG. In the case of the *BTRC* RT-PCR, we used primers specific for 5’ and 3’ sequences of the *BTRC* gene. Amplicons were loaded into the agarose gel (G-BioSciences) and visualized using Gel-Doc XR+ Imaging System (Bio-Rad).

### Co-Immunoprecipitation and Western Blot

HeLa cells were harvested and rinsed once with ice-cold 1xPBS, pH 7.2, and lysed in immunoprecipitation lysis buffer (20 mM Tris, pH 7.4, 140 mM NaCl, 10% glycerol, and 1% Triton X-100) supplemented with 1xEDTA-free complete protease inhibitor mixture (Roche) and phosphatase inhibitor cocktails-II, III (Sigma Aldrich). The cells were centrifuged at 16,000x*g* at 4°C for 30min, and the supernatants were collected. Protein concentration was quantified by modified Lowry assay (DC protein assay; Bio-Rad). The cell lysates were resolved by SDS-PAGE and transferred onto PVDF Immobilon-P membranes (Millipore). After blocking with 5% nonfat dry milk in TBS containing 0.1% Tween 20 for 1hr at room temperature, membranes were probed overnight with the appropriate primary antibodies. They were then incubated for 1h with the species-specific peroxidase-conjugated secondary antibody. Membranes were developed using the Pierce-ECL Western Blotting Substrate Kit (Thermo Scientific).

For immunoprecipitation experiments, samples were lysed and quantified as described above. Then, 3 mg of total protein was diluted with immunoprecipitation buffer to achieve a concentration of 3 mg/ml. A total of 30µl of anti-V5-magnetic beads-coupled antibody (MBL) was added to each sample and incubated for 4h at 4°C in tube rotator. Beads were then washed twice with immunoprecipitation buffer and three more times with ice cold 1xPBS. The proteins were then eluted with 40µl of 2xLaemli buffer. After a short spin, supernatants were carefully removed, and SDS-PAGE was performed. The following primary antibodies were used: anti-V5 (1:1000; Invitrogen), anti-β-catenin (1:1000; Abcam), anti-p-βcatenin (1:1000; Cell Signaling), anti-Cul1 (1:1000; Abcam), anti-SKP1 (1:1000; Cell Signaling), and anti-βactin (1:10000; Thermo Scientific).

## Supplementary Figure Legends

**Supplementary Figure 1. RNA-Seq data was obtained from BrainSpan. (A)** Schematic representation of the project workflow. Beginning with gene and isoform quantifications (processed by PsychEncode Consortium (Gandal et al., 2018; Li et al., 2018)), gene and isoform expression values were filtered based on TPM; outlier samples were removed; Surrogate Variable Analysis was performed to account for latent batch effects; temporal differential expression was performed on both datasets; WGCNA gene and isoform co-expression networks were created and analyzed. Whole exome sequencing data was obtained from Satterstrom (Satterstrom et al., 2020), filtered for LoF variants and mapped to genes and isoforms. **(B)** Initial samples were divided into distinct developmental periods as described (Kang et al., 2011; Li et al., 2018). Number of samples for each period is shown. Period P01 was omitted due to shortage of samples for the analyses.

**Supplementary Figure 2. Principal components analysis of transformed gene quantifications.** Gene expression data was transformed through regression of relevant covariates (age, brain region, gender, ethnicity, study site, and surrogate variables) to determine the appropriate number of surrogate variables (SV).16 SVs were selected for gene-level analyses.

**Supplementary Figure 3. Principal components analysis of transformed isoform quantifications.** Isoform expression data was transformed through regression of relevant covariates (age, brain region, gender, ethnicity, study site, and surrogate variables) to determine the appropriate number of surrogate variables (SV). 16 SVs were selected for isoform-level analyses.

**Supplementary Figure 4. Relative log expression analysis of transformed gene quantifications.** Gene-level relative log expression (RLE) values per sample were calculated to detect most stable relative log expression for surrogate variable selection.

**Supplementary Figure 5. Relative log expression analysis of transformed isoform quantifications.** Isoform-level relative log expression (RLE) values per sample were calculated to detect most stable relative log expression for surrogate variable selection.

**Supplementary Figure 6. Experimental validation of the isoform expression levels using independent brain samples.** The isoform expression levels of 44 splicing isoforms of 32 genes were assayed by RT-qPCR using total RNA extracted from two age and gender-matched brain samples (fetal and adult). The isoforms were selected to carry unique exonic regions. The correlation coefficient between relative expression values determined by RT-qPCR in independent samples, and those quantified from BrainSpan for the same isoforms is positive (R^2^=0.46).

**Supplementary Figure 7. ASD-LoF impacted isoforms are more highly expressed throughout development.** Isoforms that harbor ASD LoF variants are significantly more highly expressed in every brain developmental period (P < 0.0001, Wilcoxon test).

**Supplementary Figure 8**. **Gene and isoform networks have scale-free topology**. (**A**) Gene-level and (**B**) isoform-level soft-thresholding and mean connectivity, demonstrating scale-free topology of both WGCNA networks. Scale Free Topology R^2^ marked at 0.8 (red horizontal line). Both networks exhibit similar scale-free topology and have similar beta for WGCNA.

**Supplementary Figure 9**. **Gene and isoform module eigengene trajectories across brain development and associated GO functions**. Pearson correlation coefficients between all gene and isoform modules were calculated, and the highest PCC is indicated for each isoform module. The functions of isoform modules with high PCCs correspond to those of gene modules, whereas the functions of isoform modules with low PCC are generally unique. Some isoform modules also have unique eigengene trajectories across brain development (X-axes) that differ from gene module trajectories.

**Supplementary Figure 10. Contribution of cell types to the variance of BrainSpan isoform expression.** Decomposition of bulk RNA-seq BrainSpan isoform expression using fetal brain single cell RNA-seq from two studies (A) Zhong (Zhong et al., 2018) and (B) Polioudakis (Polioudakis et al., 2019) was performed using Bisque (Jew et al., 2020). Variance partition (Hoffman and Schadt, 2016) was used to investigate cell type proportions. Cell types in (B) are as follows: Excitatory deep layer 2 neurons (ExDp2), Excitatory deep layer 1 neurons (ExDp1), Maturing excitatory upper enriched neurons (ExM-U), Interneuron CGE (InCGE), Oligodendrocyte progenitor cells (OPC), Maturing excitatory neurons (ExM), Interneuron MGE (InMGE), Cycling progenitors (S phase) (PgS), Intermediate progenitors (IP), Pericytes (Per), Excitatory neurons (ExN), Cycling progenitors (PgG2M), Outer radial glia (oRG), ventricular radial glia (vRG).

**Supplementary Figure 11. Gene- and isoform-level co-expressed protein interaction networks of gM1 and iM1 modules focused on ASD risk genes.** Only edges in the top 10% of expression Pearson correlation coefficients that are also supported by gene-level protein interactions are retained. Nine genes with at least one LoF-impacted isoform are boxed.

**Supplementary Figure 12. Isoform-level co-expressed protein interaction networks of autism risk genes SCN2A, DYRK1A, DLG2 and CELF2.** Only edges in the top 10% of Pearson correlation coefficients that are also supported by gene-level protein interactions are retained.

**Supplementary Figure 13. Isoform-level co-expressed protein interaction network of *BTRC* gene.** Only edges in the top 10% of Pearson correlation coefficients that are also supported by gene-level protein interactions are retained. Non-impacted and impacted by ASD LoF *BTRC* isoforms have different partners.

## Supplementary Tables

**Supplementary Table 1.** RT**-**qPCR isoform expression validation results from two independent samples

**Supplementary Table 2.** Differential gene expression results in adjacent and Prenatal *vs* Postnatal periods

**Supplementary Table 3.** Differential isoform expression results in adjacent and Prenatal *vs* Postnatal periods

**Supplementary Table 4.** Proportions of differentially expressed genes and isoforms (summarized to gene IDs) in adjacent and Prenatal vs Postnatal periods

**Supplementary Table 5.** Gene Ontology (GO) enrichment analyses for differentially expressed genes unique to each period (red unshaded set from Fig. 1A)

**Supplementary Table 6.** Gene Ontology (GO) enrichment analyses for differentially expressed isoforms unique to each period (turquoise unshaded set from Fig. 1A)

**Supplementary Table 7.** Genes and isoforms impacted by ASD LoF mutations (extracted from Satterstrom et al. (Satterstrom et al., 2020) and processed by Variant Effect Predictor (VEP).

**Supplementary Table 8.** Gene co-expression modules identified by WGCNA.

**Supplementary Table 9.** Isoform co-expression modules identified by WGCNA.

**Supplementary Table 10.** Gene Ontology (GO) enrichment analyses for gene co-expression modules.

**Supplementary Table 11.** Gene Ontology (GO) enrichment analyses for isoform co-expression modules.

**Supplementary Table 12.** Normalized impact rate by case and control ASD LoF mutations for gene and isoform co-expression modules.

