## Supplementary Figures for "Full-length isoform transcriptome of developing human brain provides new insights into autism"

### Supplementary Figure 1

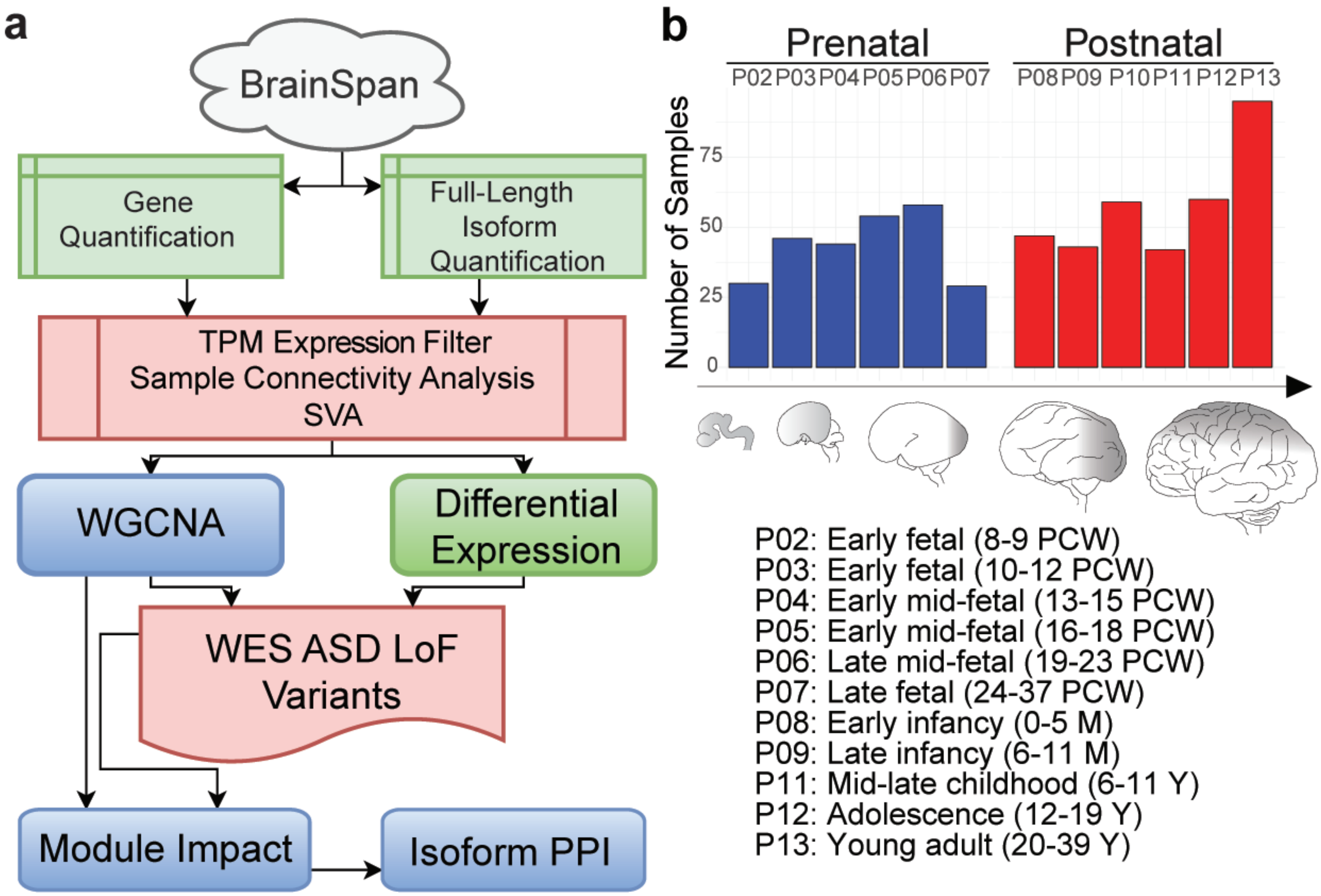


P02: Early fetal (8-9 PCW)
P03: Early fetal (10-12 PCW)
P04: Early mid-fetal (13-15 PCW)
P05: Early mid-fetal (16-18 PCW)
P06: Late mid-fetal (19-23 PCW)
P07: Late fetal (24-37 PCW)
P08: Early infancy (0-5 M)
P09: Late infancy (6-11 M)
P11: Mid-late childhood (6-11 Y)
P12: Adolescence (12-19 Y)
P13: Young adult (20-39 Y)

### Supplementary Figure 2

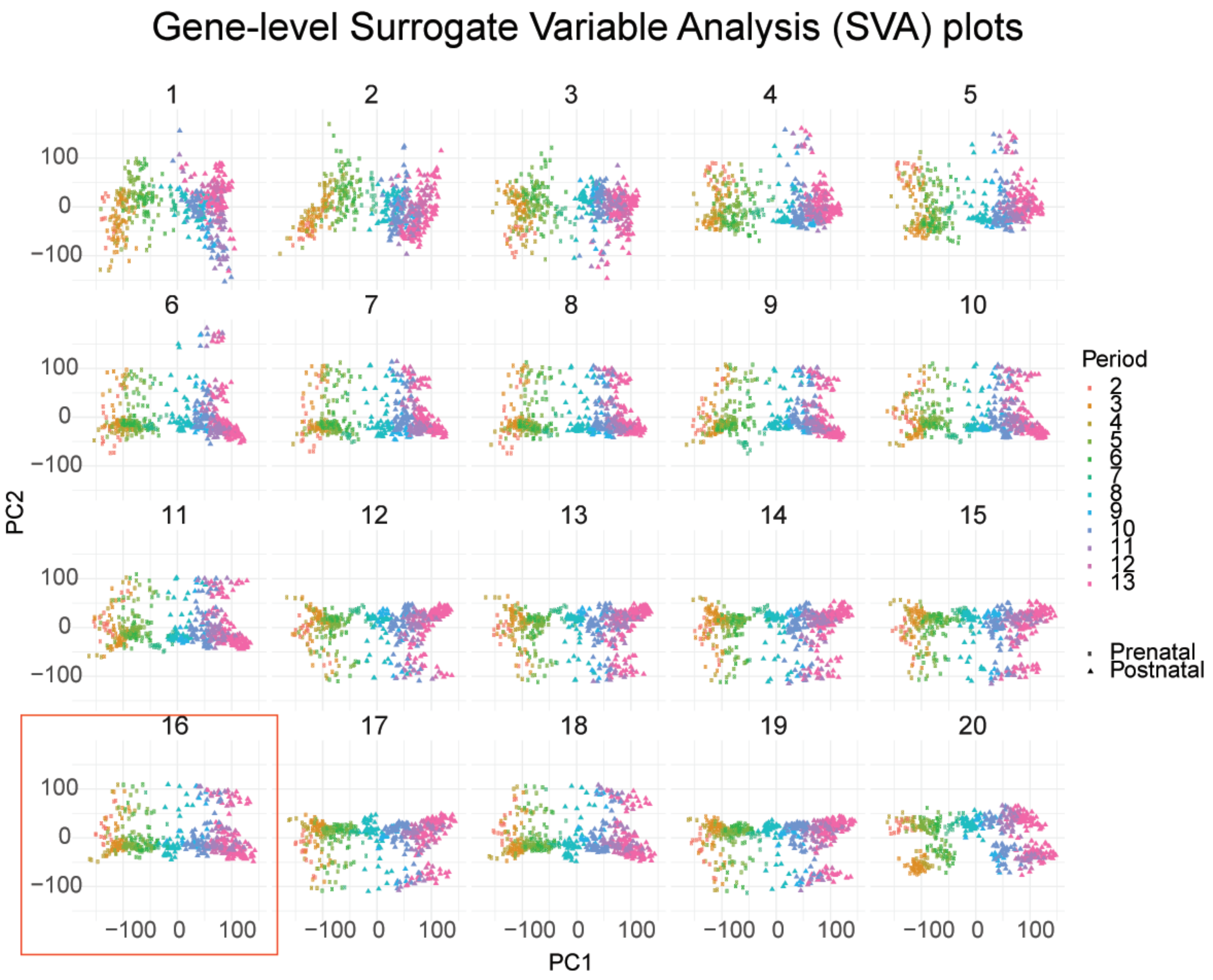

### Supplementary Figure 3

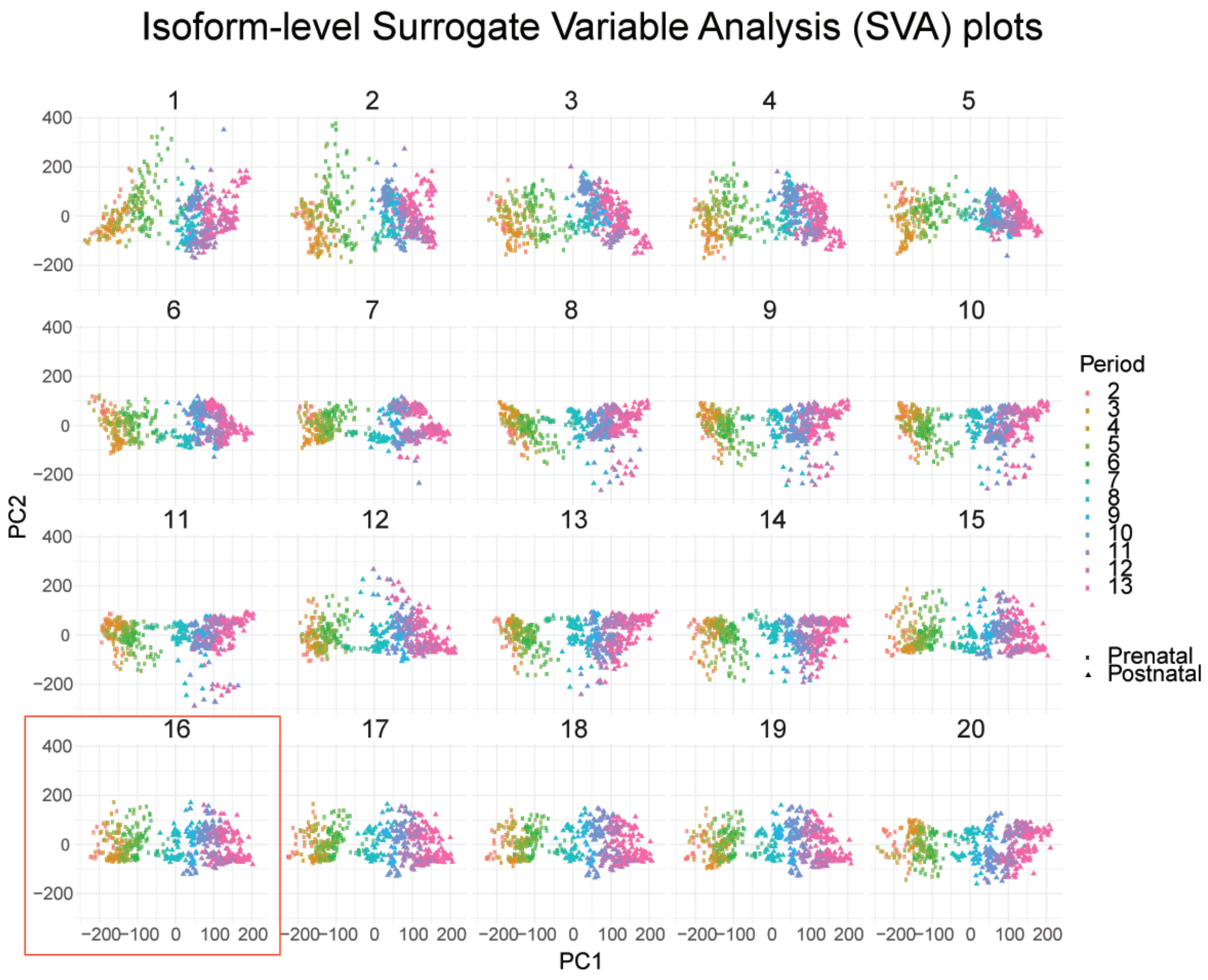

### Supplementary Figure 4

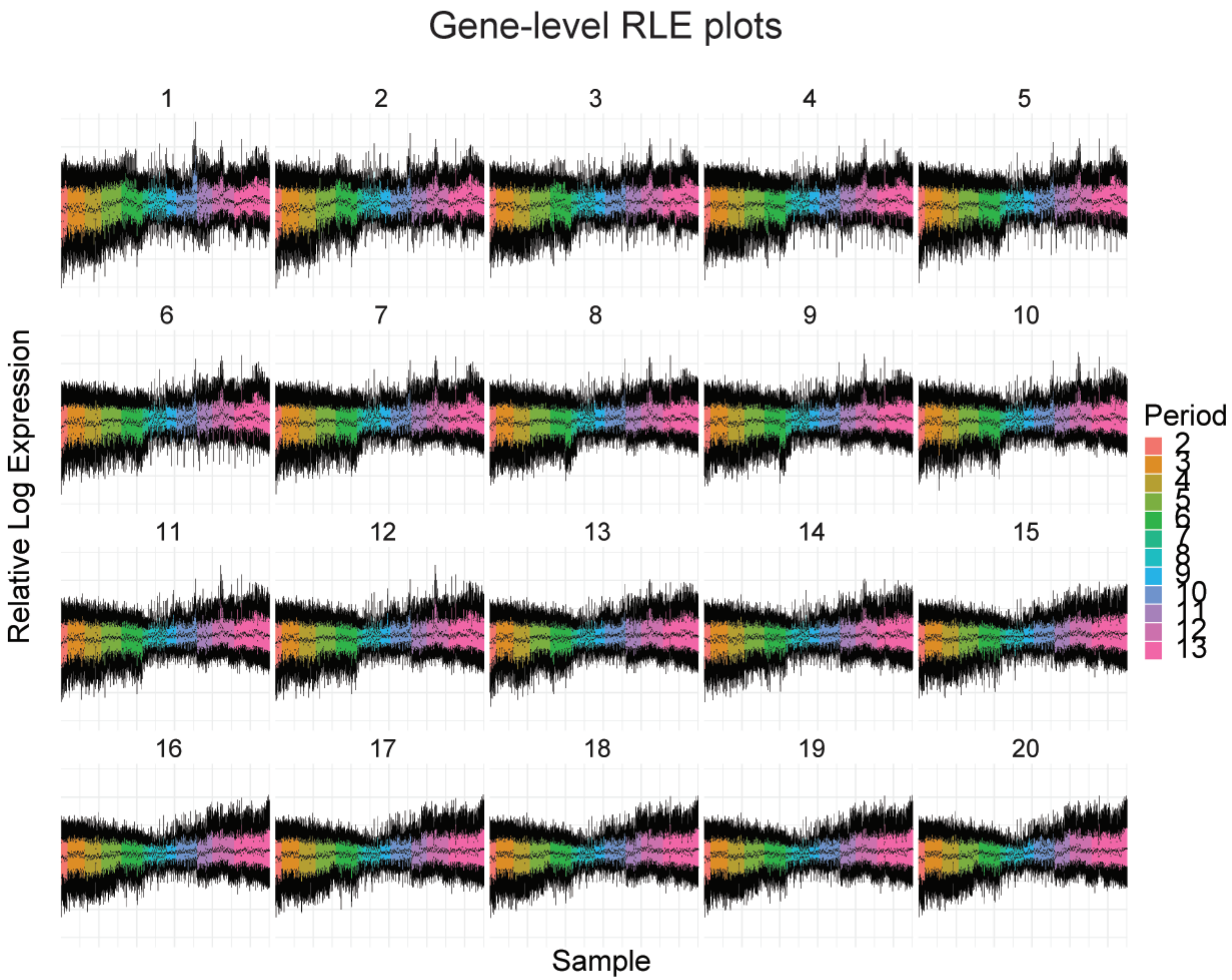

### Supplementary Figure 5

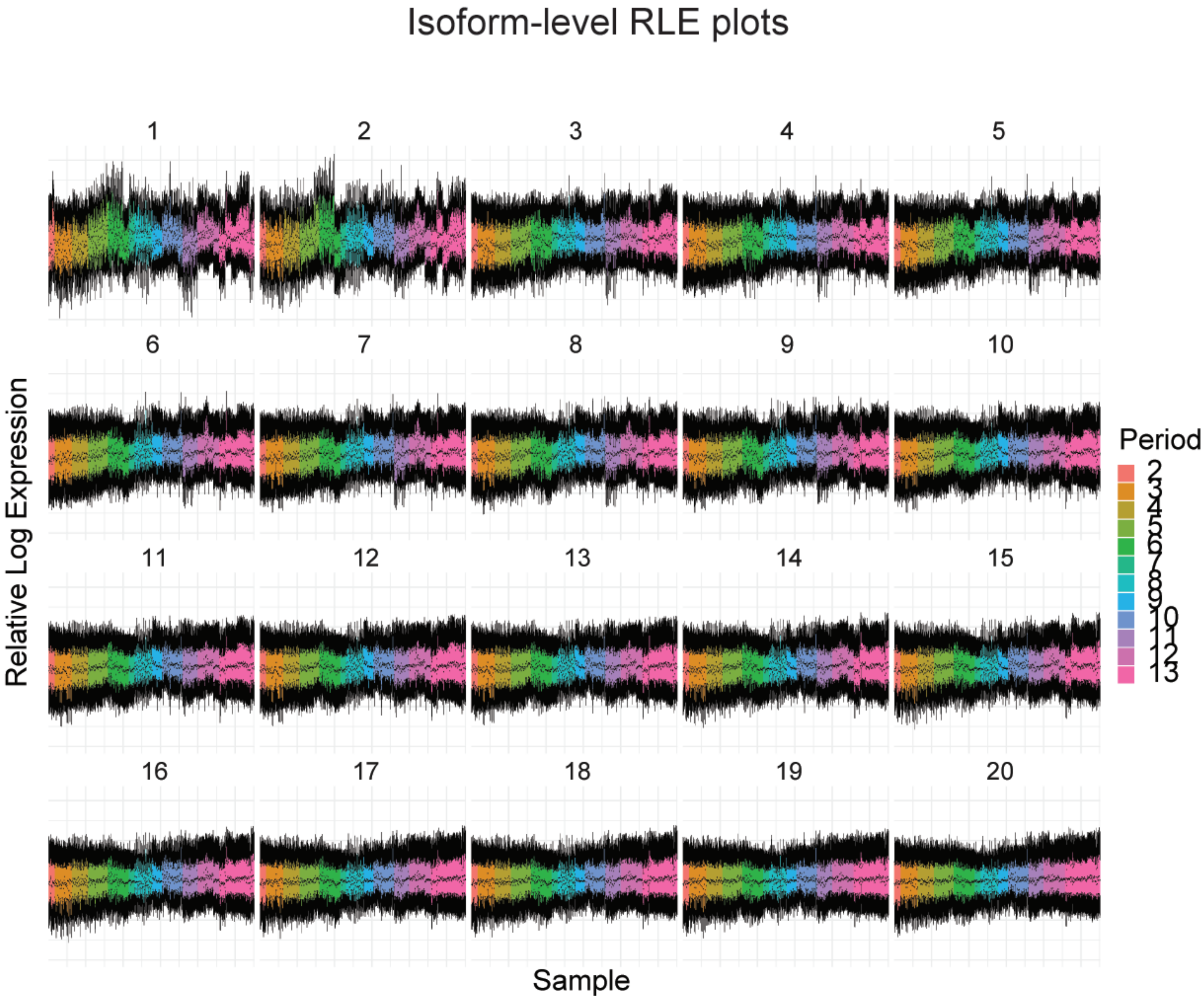

Supplementary Figure 6

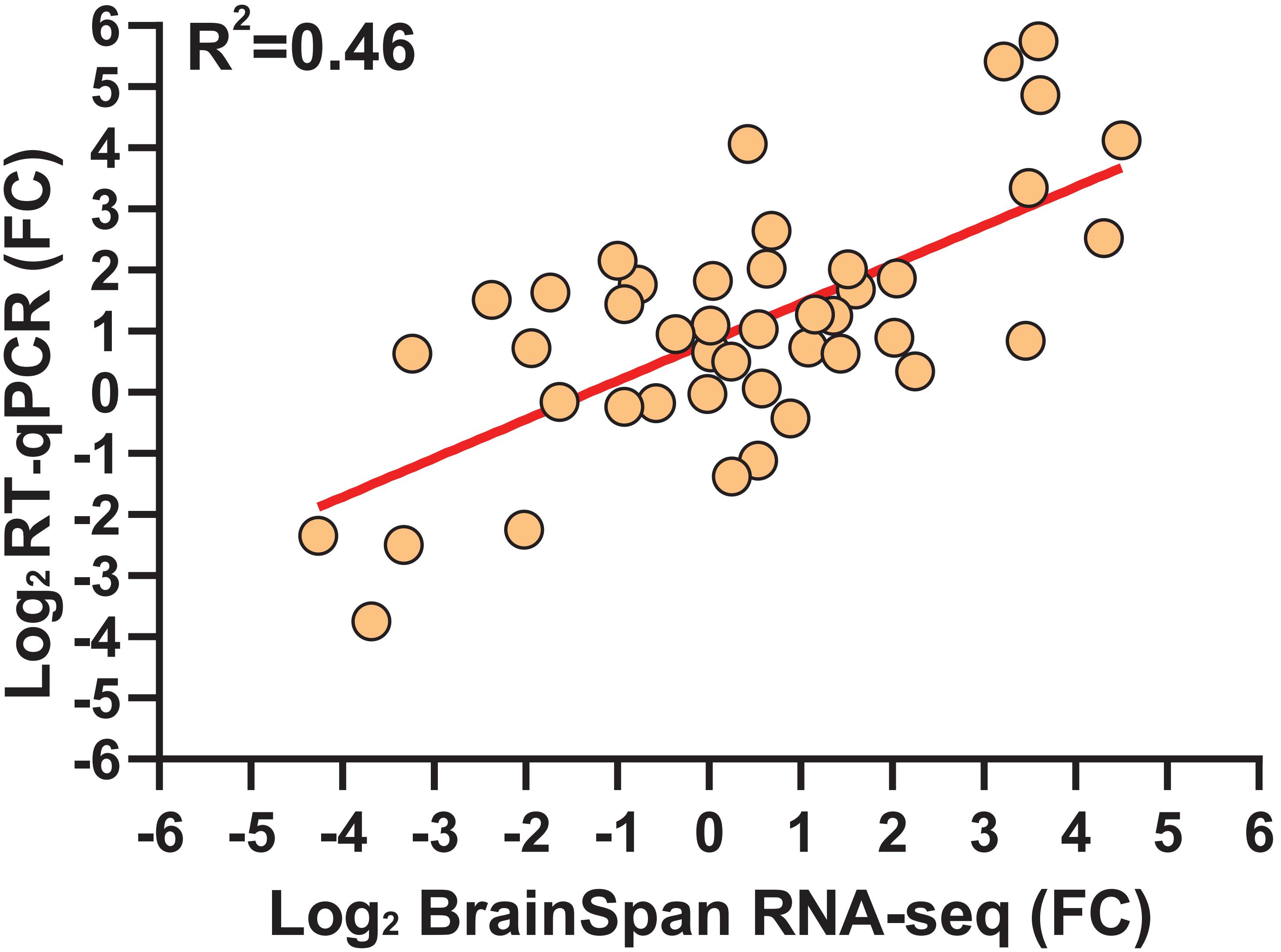

### Supplementary Figure 7

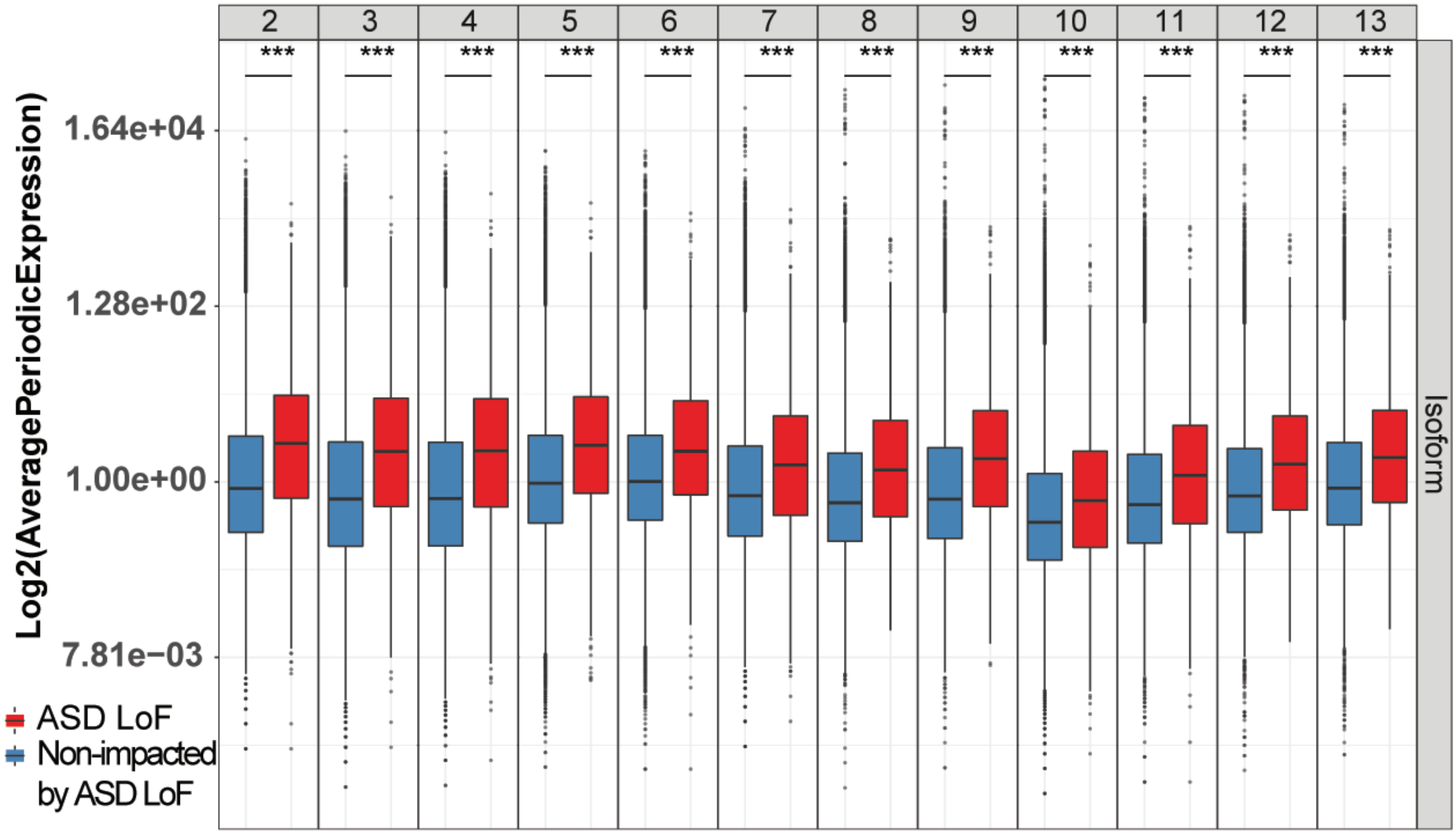

### Supplementary Figure 8

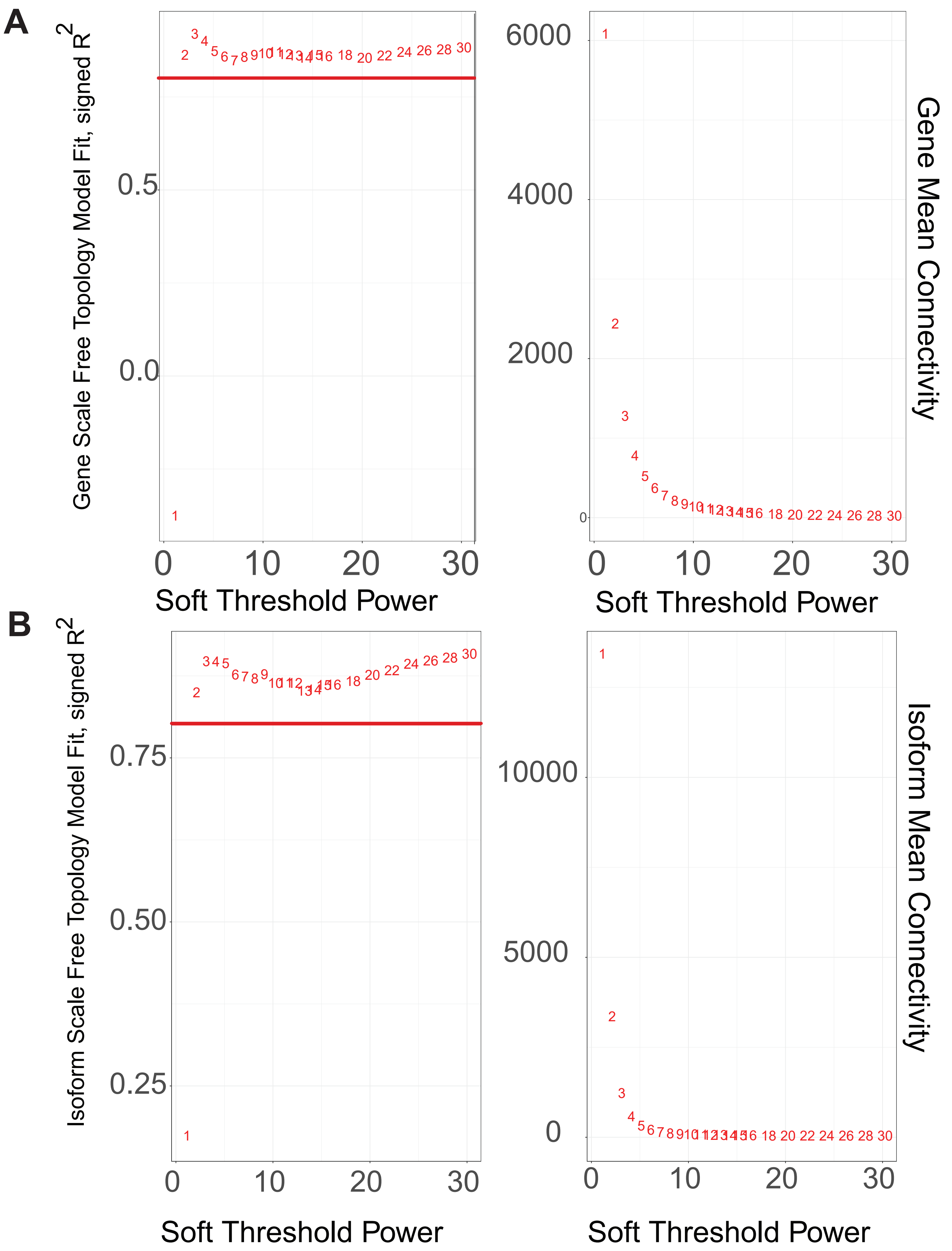

### Supplementary Figure 9

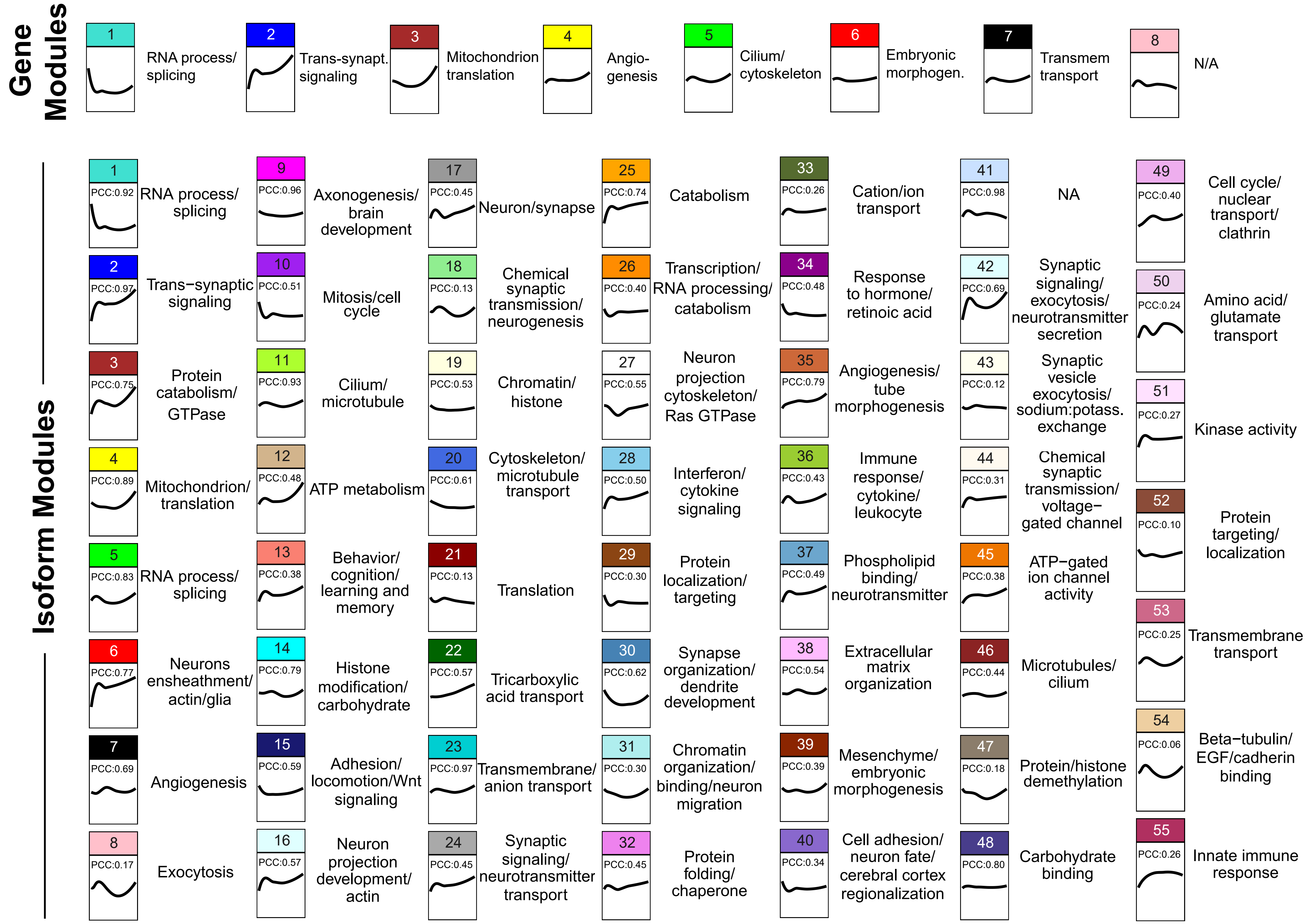

### Supplementary Figure 10

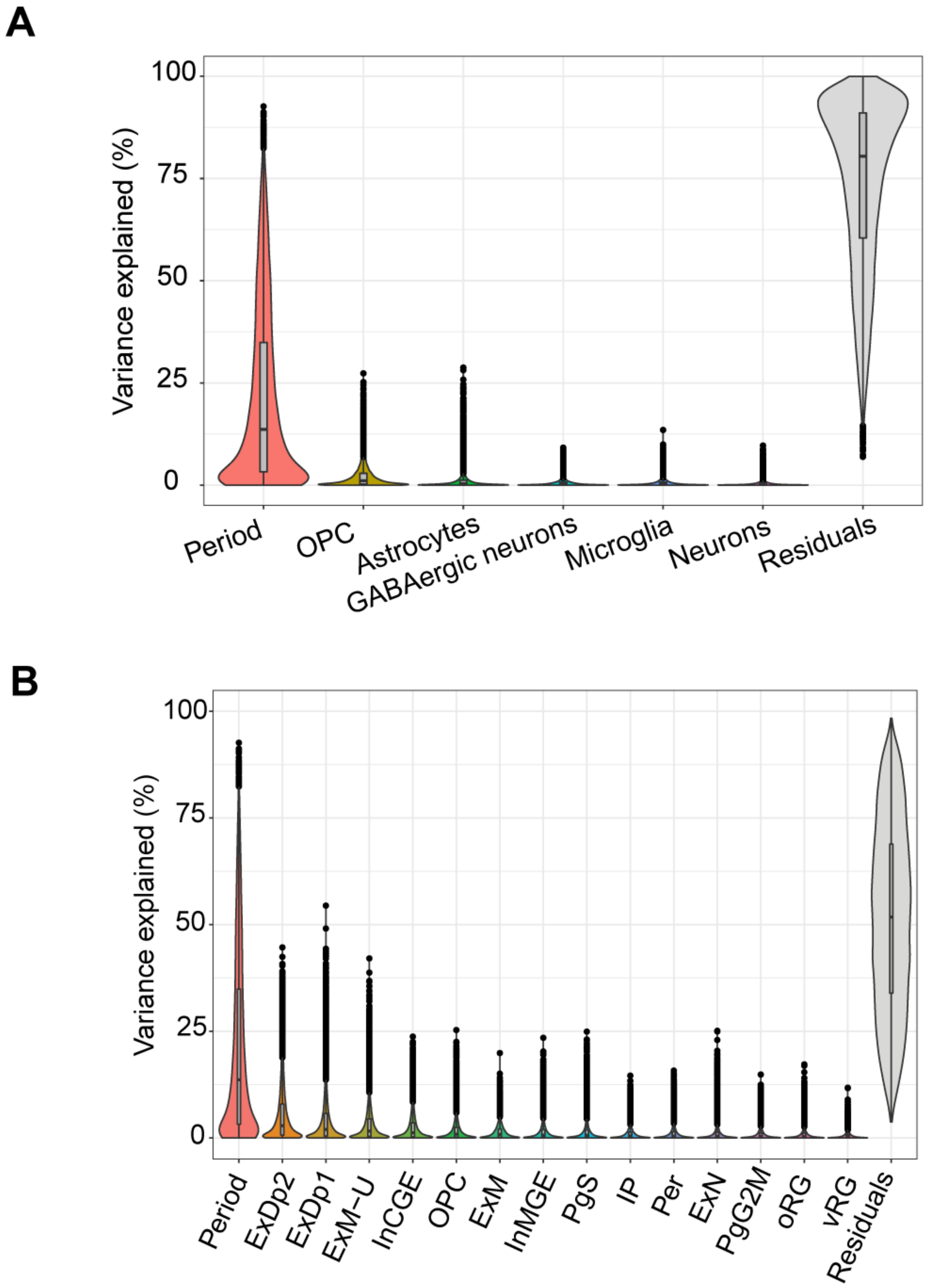

### Supplementary Figure 11

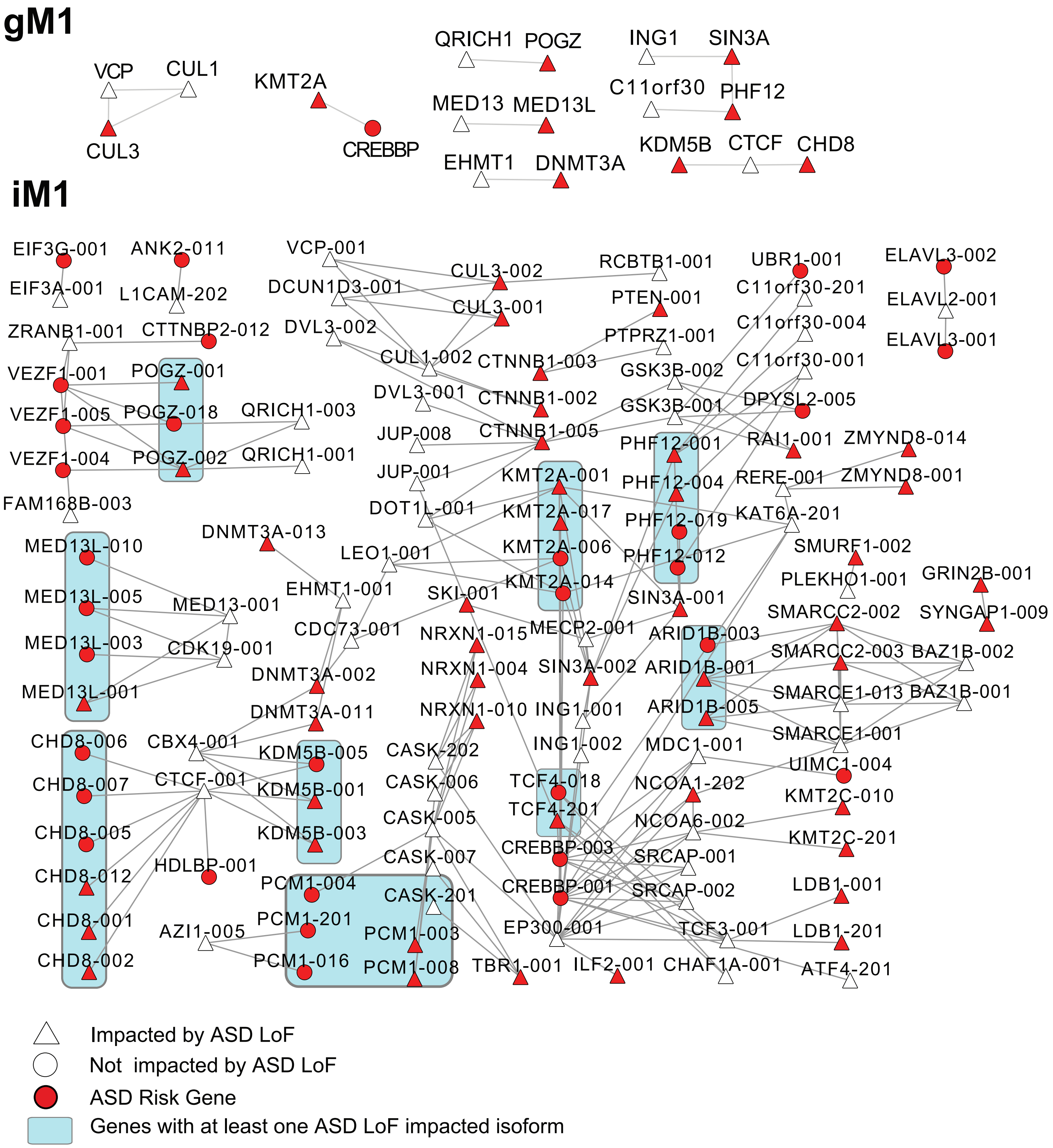

### Supplementary Figure 12

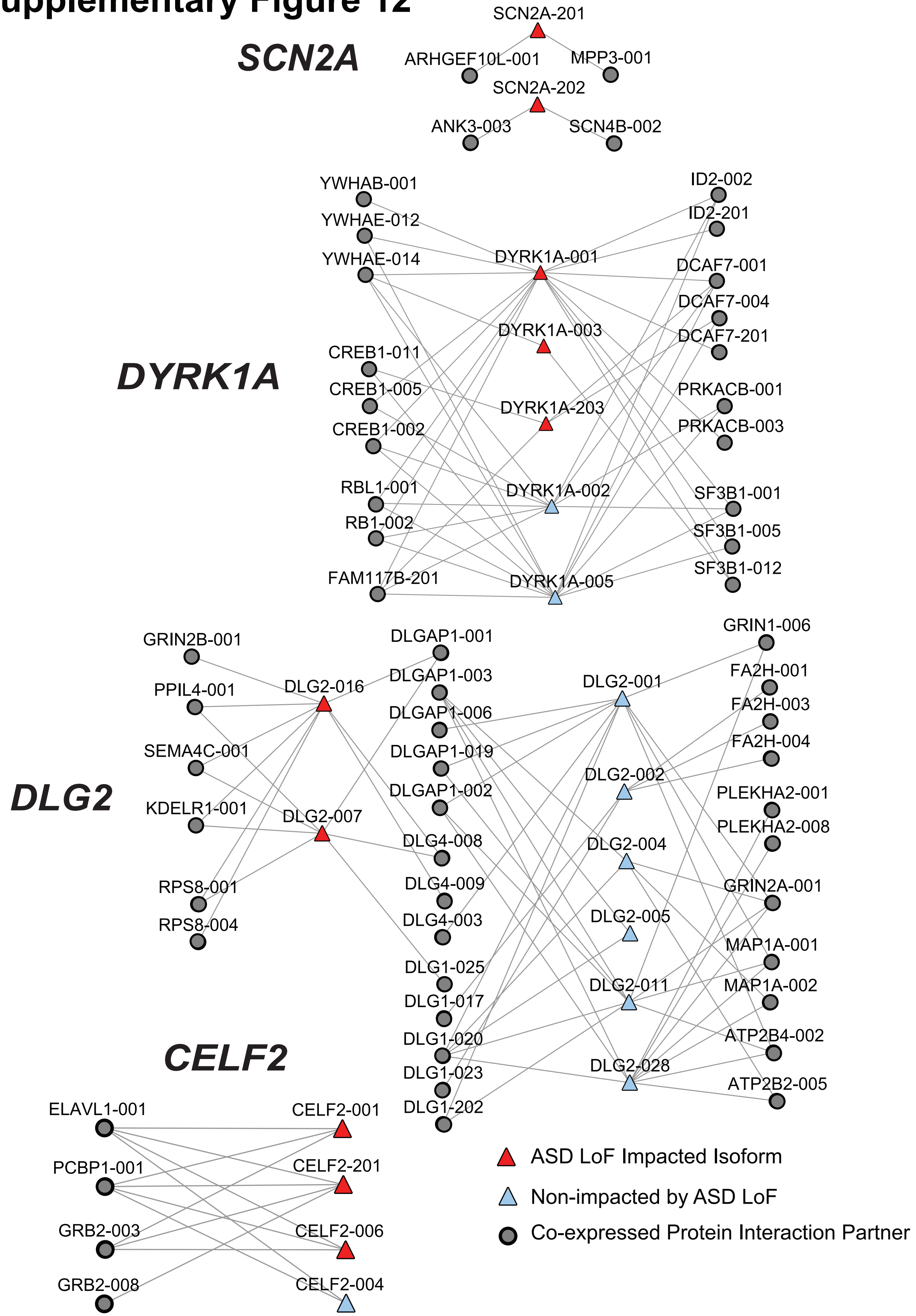

#### Supplementary Figure 13

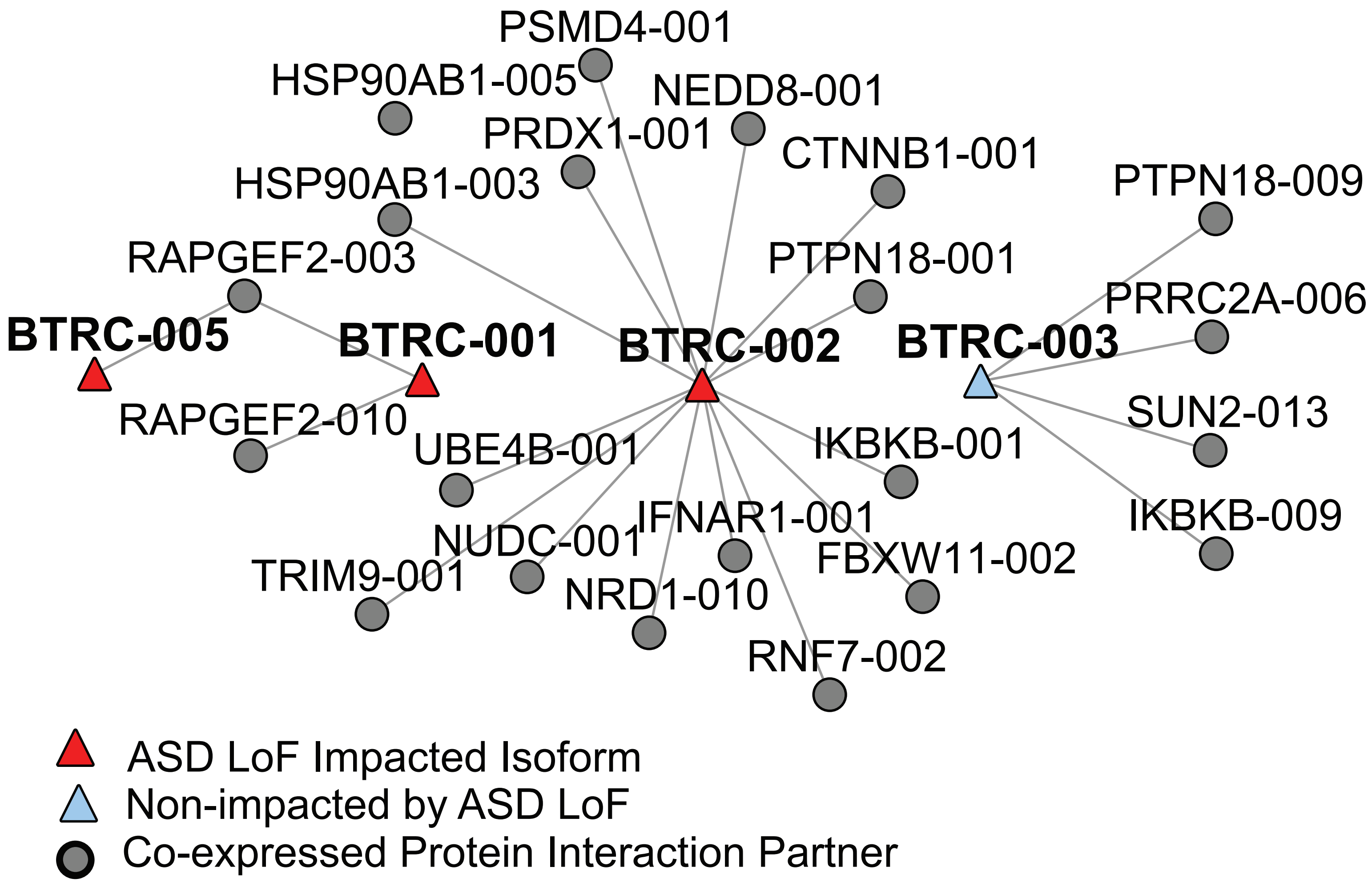
